# Disease stage-specific pathogenicity of CD138 (syndecan 1)-expressing T cells in systemic lupus erythematosus

**DOI:** 10.1101/2020.03.25.008995

**Authors:** Lunhua Liu, Kazuyo Takeda, Mustafa Akkoyunlu

## Abstract

**Objective:** To identify and characterize CD138 (syndecan 1)-expressing T cells in SLE-prone mice.

**Methods:** We characterized CD138-expressing T cells in MRL/Lpr mice by flow cytometry assay and by gene analysis. Functional properties of TCRβ+CD138+ cells were assessed either by activating through TCR or by co-incubating with purified B cells in the presence of auto-antigens. Purified TCRβ+CD138+ cells were adoptively transferred into MRL/Lpr mice and lupus disease was assessed by measuring serum auto-antibodies, proteinuria and by histopathological evaluation of kidney.

**Results:** We found that the frequency of TCRβ+CD138+ cells was significantly higher in MRL/Lpr mice than in wild-type MRL mice (p < 0.01), and the increase in their numbers correlated with disease severity. Majority of the TCRβ+CD138+ cells were CD4 and CD8 double-negative and 20% were CD4. Compared to TCRβ+CD138− cells, TCRβ+CD138+ cells exhibited central memory phenotype with reduced ability to proliferate, and produce the cytokines IFNγ and IL-17. When co-cultured with B cells, the ability of TCRβ+CD138+ cells to promote plasma cell formation and autoreactive antibody production was dependent on the presence of auto-antigens and CD4 co-receptor expression. Surprisingly, adoptively transferred TCRβ+CD138+ cells slowed down the disease progression in young MRL/Lpr mice but had the opposite effect when DNA was co-administered or when TCRβ+CD138+ cells were transferred into older MRL/Lpr mice with established disease.

**Conclusion:** Here, we provided evidence for the pathogenic role of CD138-expressing T cells when auto-antigens are exposed to immune system. Thus, monitoring the changes in TCRβ+CD138+ cell-frequency may serve as a tool to assess SLE severity. Moreover, CD138-expressing T cells may be targeted to alleviate lupus progression.

## INTRODUCTION

Systemic lupus erythematosus (SLE) is an autoimmune disease characterized by the production of autoantibodies and inflammatory cell infiltration in multiple tissues [1]. T cells are critical for SLE pathogenesis as they, not only provide help to autoreactive B cells, but also infiltrate and damage the target organs, such as the skin, joints, brain, lung, heart and kidneys [2]. T cells from SLE patients and lupus-prone mice have expanded T helper 1 (Th1), Th17, follicular and extrafollicular helper T (Tfh and eTfh) cell subsets, which contributes to the inflammation and autoreactive antibody production through increased IFNγ, TNFα, IL-6, IL-17, and IL-21 secretion [3–7]. Diminished populations of T regulatory cells (Treg), T follicular regulatory (Tfr) cells and cytotoxic CD8+ T cells were also thought to be contributing to the pathogenesis of SLE [8–11]. In addition, increased circulating TCRαβ+CD4-CD8− double-negative (dn) T cells and renal accumulation of the minor γδT population have been associated with autoantibody production and lupus nephritis [12]. Besides the phenotypic and functional alterations in effector T cells, terminally differentiated memory T cells may also be contributing to the tissue damage as these cells accumulate in SLE patients with high disease activity [13–15].

Syndecans, type I transmembrane heparan sulphate proteoglycans (HSPG), regulate diverse biological processes, such as tissue wound repair, angiogenesis, epithelial-mesenchymal transformation, and inflammation, by modifying the local concentration, stability and accessibility of extracellular matrix components, cytokines, chemokines and growth factors [16]. Syndecan family consists of four distinct members that are mostly expressed on epithelial, endothelial, neural or fibroblastic cells, but they are also detected on haemopoietic cells [17, 18]. Syndecan 2 and syndecan 4 are up-regulated upon CD4+ T cell activation and act as inhibitors by promoting T cell receptor (TCR) clearance or by activating tyrosine phosphatase CD148 [19, 20]. Syndecan 1 (CD138) is commonly used as a marker to identify plasmablasts and plasma cells [21]. Recently, CD138 has been suggested to be a a marker to distinguish IL-17 producing natural killer T17 (NKT17) cells from NKT1 cells, based on its selective expression on NKT17 cells but not on NKT1 cells [22, 23]. Moreover, CD138+ T cells were observed accumulated in gut epithelia of aged C3H wild type as well as in the spleen and lymph nodes of FasL loss-of-function C3H gld mice [24]. Similarly, CD138-expressing T cells were detected in spleen and lymph nodes of lupusprone, μMT/lpr mouse but these cells were only present in the lymph nodes, and not in the spleen, of another lupus-prone strain, B6/lpr mouse [25]. Thus, accumulating evidence indicate the existence of CD138+ T cells in both healthy and diseased mice. However, the characteristics and pathologic roles of CD138+ T cells in lupus disease remain to be elucidated.

Here, we detected the presence of TCRβ+CD138+ cells in various organs of the lupus-prone, MRL/Lpr mice. The numbers of TCRβ+CD138+ cells increased as the disease progressed. We also identified CD4+ T cells among the TCRβ+CD138− population as an important source of TCRβ+CD138+ cells. These accumulating TCRβ+CD138+ cells manifested mostly central memory phenotype (Tcm) and promoted lupus disease progression only when autoantigens were present, although they exhibited slower activation kinetics, less proliferation and diminished cytokine production after stimulation with anti-CD3/CD28 antibodies.

## METHODS

### Mice

MRL/MpJ-FASLPR/J (referred to as MRL/Lpr throughout the manuscript), MRL/MpJ (referred to as MRL throughout the manuscript) mice, and C57BL/6 mice were purchased from The Jackson Laboratory (Bar Harbor, ME). Balb/c mice were purchased from Charles River Laboratories (Wilmington, MA). Only age-matched female mice were used for experiments. All mice were bred and maintained under specific pathogen-free conditions in the animal facility of US Food and Drug Administration (FDA), Center for Biologics Evaluation and Research (CBER) Veterinary Services. The breeding and use of animals were approved by the US FDA, CBER Institutional Animal Care and Use Committee (permit numbers 2002-37 and 2017-47).

### Detection of anti-dsDNA and SM antibodies in sera

Serum anti-dsDNA and anti-Smith antigen (SM) antibodies were measured by ELISA as described previously [26]. Briefly, calf thymic DNA (Sigma, St. Louis, MO) or SM (Immunovision, Springdale, AR) were coated on 96-well microtiter plates (Dynatech Immulon 4 HBX; Dynatech Labs., Chantilly, VA) at 0.5 μg/ml with 0.1 M of carbonate-bicarbonate buffer (pH 9.6) overnight at 4°C. Plates were blocked for 30 minutes at room temperature in 5% BSA in PBS, then washed with 0.05% Tween-20 in PBS. Diluted serum samples were added to wells in triplicates and incubated at 37°C for 2 hours. Plates were washed with 0.05% Tween-20 in PBS and further incubated with HRP-conjugated goat antibodies directed against mouse IgG (Southern Biotech, Birmingham, AL) for 1 hour at room temperature. Finally, plates were washed with 0.05% Tween-20 in PBS and measured at 405 nm absorbance after developing with ABTS solution (Invitrogen, Carlsbad, CA). Antibody titers were recorded as the last titration corresponding to the OD that is twice the mean OD of blank wells.

### Flow cytometry

Single cell suspensions of spleen, bone marrow, lymph nodes, and thymus were obtained by mechanic dissociation of tissue through a 40 μm cell strainer. The dissociated cells were filtered through a 100 μm cell strainer. Red blood cells were then lysed using ACK lysing buffer (Lonza, Wallersville, MD). In addition, mouse blood leukocytes were collected by lysing red blood cells with ACK lysing buffer and centrifugation at 300 × g for 5 minutes. Cells were stained with fluorescent-conjugated anti-mouse antibodies after blocking CD16/CD32 with Fc Block (BD Biosciences, San Jose, CA). For intracellular staining, Brefeldin A (BD Biosciences)-treated cells were stained with the surface markers and LIVE/DEAD™ Fixable Near-IR Dead cell kit (Fisher Thermo,Waltham, MA) before fixation, permeabilization and intracellular staining as per manufactures instructions (BD Biosciences). The following antibodies were used in flow cytometry analysis: Pacific blue anti-CD19, BV421 anti-CD19, BV421 anti-TCRβ, APC anti-CD138, APC anti-TCRβ, BV605 anti-CD3, FITC anti-CD3, Percp Cy5.5 anti-CD44, FITC anti-602L, PE-Cy7 anti-PD-1, APC anti-CXCR5, Percp Cy5.5 anti-B220, PE-Cy7 anti-CD8, PE anti-CD21, PE anti-CD22, BV421anti-CD23, Alexa647 anti-CD40, FITC anti-CD80, FITC anti-CD86, Percp Cy5.5 anti-CD25, FITC anti-CD69, APC anti-CD95, Percp Cy5.5 anti-IL17, Percp Cy5.5 anti-CCR7, FITC anti-FOXP3 (all purchased from BioLegend, San Diego, CA). PE-anti-CD138 was purchased from BD biosciences. In addition, FITC anti-BCMA, PE anti-TACI, FITC anti-IFNγ, Annexin V, (R&D system, Minneapolis, MN), ATTO 488 anti-BAFFR (Enzo life Science Inc., Farmingdale, NY), CellTrace^™^ CFSE Cell Proliferation Kit and Qdot605 anti-CD4 antibody (Thermo Fisher). Stained cells were acquired using LSR II flow cytometer (BD Biosciences) and data were analyzed using FlowJo (Tree Star, Ashland, OR) version 10.1 for PC.

### Quantitative Real-Time PCR

Total RNA was extracted from flow cytometry-sorted cells using the RNeasy Mini kit (Qiagen, Germantown, MD). Two hundred nanograms of total RNA were reverse-transcribed into cDNA using random hexamers with the Taqman Reverse transcription kit (Invitrogen). The expression of targeted genes and GAPDH were determined using Taqman Gene Expression assays and CFX96 Touch Real-Time System (BioRad, Hercules, CA). Relative expression values were determined by the 2-ΔCt method where samples were normalized to GAPDH gene expression.

### T cell isolation and adoptive transfer experiments

Splenic T cells from MRL/Lpr mice were purified with Dynabeads^™^ FlowComp^™^ Mouse Pan T (CD90.2) Kit and dissocated from beads as per manufacture’s instructions (ThermoFisher). Purified T cells were staind with PE-conjugated anti-CD138 antibody, and TCRβ+CD138+ and TCRβ+CD138− cells were further separated with anti-PE magnetic MicroBeads (Miltenyi Biotec, Auburn, CA). After three washes with PBS, cells were suspended in PBS and 1 × 10^7^ cells in 100 μl were i.v. injected into recipient mice. TCRβ+CD4+CD138− cells were isolated from TCRβ+CD138− cells using the CD4 (L3T4) MicroBeads (Miltenyi Biotec), and unbound cells were identified as TCRβ+CD8+CD138− cells.

### Co-culture of B cells with T cells

Splenic B cells were isolated from 5 or 12 weeks old MRL/Lpr mice using B Cell Isolation Kit (Miltenyi Biotec) and stained with CSFE before co-culturing with purified TCRβ+CD138+ or TCRβ+CD138− cells in the presence of anti-CD3/CD28 antibodies (BD Biosciences), phorbol 12-myristate 13-acetate (PMA)/ionomycin or autoantigens (1 μg/ml of DNA or SM (Immunovision)). DNA was isolated from MRL/Lpr splenocytes by hyperthemo treatment at 42°C for 4 hours. After 3-4 days of incubation, cells were analyzed for CFSE dilution by flow cytometry. In other assays, after 10 days of culture, culture supernatants were analyzed for antibody production as well as IL-2 and IFNγ secretion by ELISA (R&D Systems). In CD4 blocking expriments, Ultra-LEAF^™^ purified CD4 antibody (clone GK1.5, Biolegend) or control rat IgG (Sigma-Aldrich) were added to T and B cell co-cultures. In some co-culture experiments, T and B cells were incubated either as mixed or separated with 0.4 μm pore sized polyester Corning Transwell® membrane insert (Sigma-Aldrich).

### Pristane-induced lupus model

Nine weeks old female Balb/c or C57BL/6 mice received a single i.p. injection of 0.5 ml pristane (Sigma-Aldrich) or 0.5 ml of sterile PBS. Four months later, sera were collected and spleens were harvested. Sera were analyzed for auto-antibodies by ELISA, and splenocytes were subjected to flow cytometry for the presence of TCRβ+CD138+ cells.

### Evaluation of disease progression and histopathological assessment of the kidneys

Proteinuria was measured using Fisherbrand^™^ Urine Reagent Strips (Fisher scientific, Hampton, NH) and scored on a scale of 0-5 (0, none; 1, trace; 2, 30 mg/dl; 3, 100 mg/dl; 4, 300 mg/dl; and 5, ≥2000 mg/dl).

### Histopathological analysis

Mouse kidneys were fixed in 10% buffered formalin overnight, processed, and embedded in paraffin. Sections were processed as previously described [26]. Hematoxylin and eosin (H&E), Masson trichrome, periodic acid-Schiff (PAS) and acid methenamine silver (PAM) stainings were performed. Stained slides were scanned by Nanozoomer XR (Hamamatsu corporation, Japan) and data was store as ndpi files for analysis. Overall severity, glomerular sclerosis, inflammatory cell accumulation, and interstitial fibrosis were evaluated and scored semiquantatively between 0-3 (0=within normal limits, 1=mild pathology, 2=moderate pathology and 3=severe pathology). Average scores were analyzed by GraphPad Prism (GraphPad Software, San Diego, CA) software.

### Statistical analysis

Data from groups were compared using GraphPad Prism software and nonparametric testing was performed by the Mann-Whitney rank sum test for two groups and by Kruskal-Wallis two-way ANOVA on ranks for three or more groups.

## RESULTS

### Frequency of CD138-expressing TCRβ+ cells increases parallel to disease progression in MRL/Lpr mice

The presence of CD138-expressing αβ T cells has been reported in Fas and FasL mutant mouse strains (C3H gld, μMT/lpr and B6/lpr mice) manifesting lupus-like disease, but the physiological significance of CD138-expressing T cells in lupus pathogenesis remains unexplored [24, 25]. Here, we investigated the immunopathological role of CD138-expressing T cells in MRL/Lpr mouse, a widely used lupus-prone strain [27]. First, we found a large fraction of CD138 expressing cells among CD19-TCRβ+ gated splenic population in MRL/Lpr mice (6 weeks old), even before the onset of autoimmune manifestations (Figure 1A, Supplemental Figure 1A). Measurement of CD138 mRNA in qRT-PCR assay on sorted TCRβ+CD138+ cells confirmed the expression of CD138 (Supplemental Figure 1B). However, the frequency of TCRβ+CD138+ cells in the spleens of age-matched Balb/c, C57BL/6 as well as the parental MRL mice remained negligible (Figure 1A). Thus, the expression of CD138 on T cells appears to be uniquely associated with Fas signaling deficiency [24, 25]. Second, we investigated a possible correlation between TCRβ+CD138+ cells and the progression of disease in MRL/Lpr mice and found that the frequency of TCRβ+CD138+ cells progressively increased with the age of mice (Figure 1B). Moreover, the increase in TCRβ+CD138+ cell population was also detected in pristine-injected Balb/c and C57BL/6 mice (Supplemental Figure 1C and D), which also develop lupus-like autoimmune symptoms [27]. It is important to note that in B6/lpr mice, TCRβ+CD138+ cells were previously reported to be confined to the lymph nodes [25], however, we found high percentage of these cells also in the thymus, spleen, lymph nodes and blood of MRL/Lpr mice (Figure 1C). The increase in blood reached to over 30% of blood lymphocytes in 14-week old mice, suggesting extensive circulation of TCRβ+CD138+ in MRL/Lpr mice when they develop severe lupus disease.

**Figure 1.**
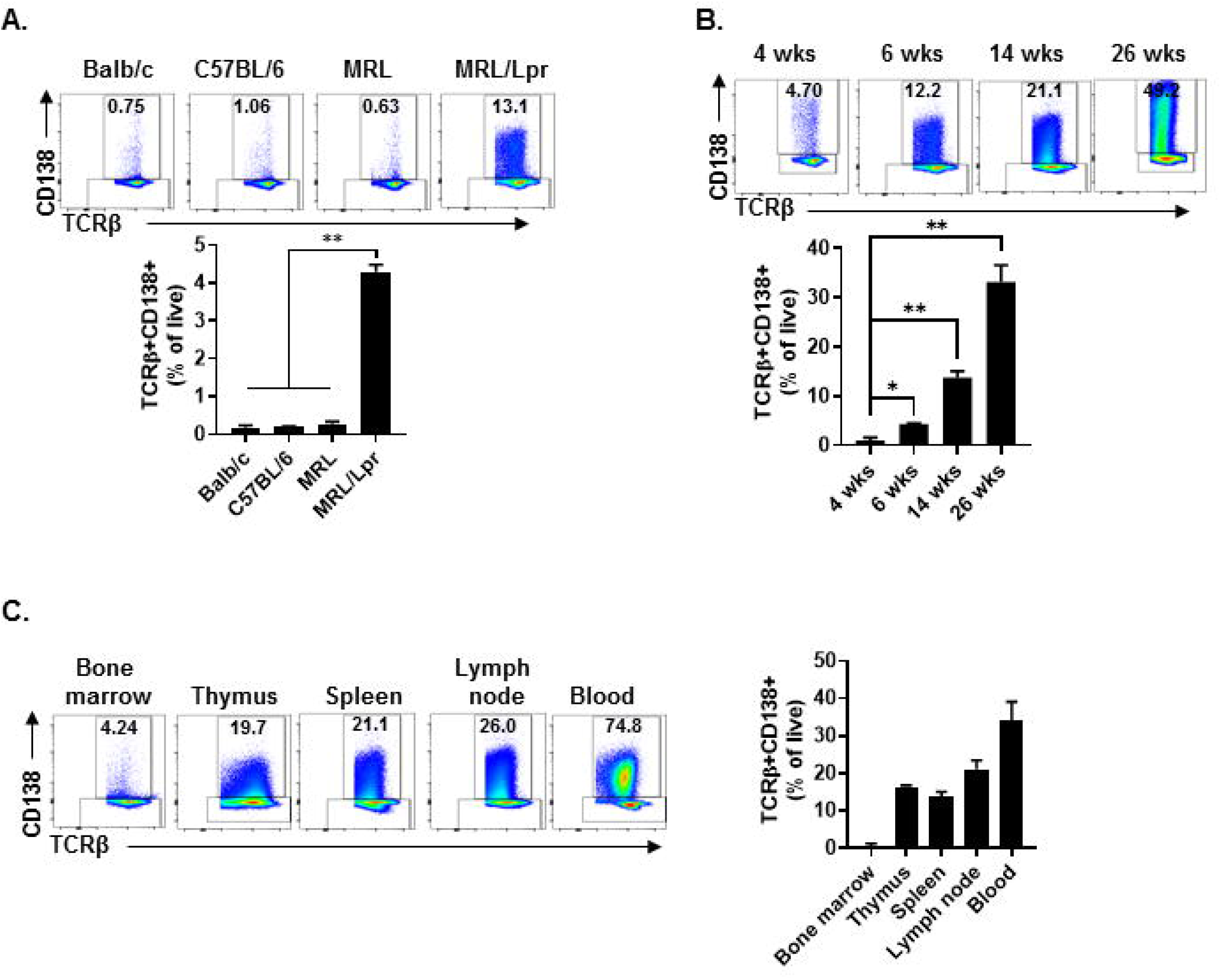
TCRβ+CD138+ cells populate the spleen, lymph nodes, thymus and blood of MRL/Lpr mice, and their numbers increase with age. In each flow cytometry experiment TCRβ+CD138+ cells were quantified after gating out dead cells and CD19+ B cells. **(A)** Splenic TCRβ+CD138+ cells from 6 weeks old Balb/c, C57BL/6, MRL and MRL/Lpr mice were quantified. Representative pseudocolor plots from each mouse strains are shown. Mean percentages ± SD of 5 mice from two independent experiments are plotted. **p<0.01. **(B)** Splenic TCRβ+CD138+ cells from 4, 6, 14 and 26 weeks old MRL/Lpr mice were quantified. Representative pseudocolor plot from each time point is shown. Mean percentages ± SD of 5 to 8 mice from three separate experiments are plotted. *p<0.05, **p<0.01 **(C)** TCRβ+CD138+ cells in bone marrow, thymus, spleen, lymph nodes and blood of 14 weeks old MRL/Lpr mice were quantified. Representative pseudocolor plots from each organ are shown. Mean percentages ± SD of 5 mice from two independent experiments are plotted.

### TCRβ+CD138+ cells derive from CD138-CD4+ T cells

Next, we further characterized the MRL/Lpr mice TCRβ+CD138+ cells by assessing the expression of surface markers and the genes associated with B and T cell lineages. As shown previously in μMT/lpr and B6/lpr mice [24, 25], most of TCRβ+CD138+ cells also expressed B220, while only about 5% of TCRβ+CD138− did (Supplemental Figure 2A). There was no apparent difference in the expression of other B cell-related surface proteins BAFFR, BCMA, IgM, CD21, CD23, CD40, CD80, or CD86 as well as mRNA for *CD19*, *IGHM, Myc*, *tnfrsf13c*, and *tnfrsf13b* between TCRβ+CD138+ and TCRβ+CD138− cells (Supplemental Figure 2B to D). In addition, the expression levels of transcription factors (*Pax5*, *PU1*, *Irf4*, and *Xbp1*) associated with B and plasma cells were comparable between TCRβ+CD138+ and TCRβ+CD138− cells, although *Bcl-6* was higher and *Prdm1* lower in CD138-expressing cells (Supplemental Figure 2E). CD138 expression did not affect the expression of the T cell marker CD3 on TCRβ+ cells as both TCRβ+CD138+ and TCRβ+CD138− were CD3+ (Figure 2A and B). In μMT/lpr and B6/lpr mice, all TCRβ+CD138+ cells were reported to be negative for CD4 and CD8 [24, 25]. Although the majority of TCRβ+CD138+ cells were negative for CD4 and CD8 in MRL/Lpr mice also, approximately 20% of TCRβ+CD138+ were CD4+, while only ~2% were CD8+ (Figure 2A and B). Further confirming the association of TCRβ+CD138+ cells with T-cell but not with B-cell lineage, we measured comparable levels of *GATA3* and *Tbet* expression in TCRβ+CD138+ and TCRβ+CD138− cells, the transcription factors associated with Th2 and Th1 subsets [28], respectively (Figure 2C). Interestingly, CD138− cells expressed the Treg cell transcription factor *Foxp3* but not CD138-expressing cells (Figure 2C). To assess whether CD138 expression changed over time in incubated TCRβ+ T cells, next we assessed the changes in the percentage of CD138+ cells in cultured TCRβ+CD138+ and TCRβ+CD138− cells *in vitro*. As shown in Figure 2D, CD138 levels remained high on TCRβ+CD138+ cells throughout the 7-day culture period. By contrast, there was a gradual increase in the frequency of CD138+ cells among the cultured TCRβ+CD138− cells from day 1 to day 3 (Figure 2D). We next sought to determine whether CD138+ cells emerged from TCRβ+CD4+CD138− or TCRβ+CD8+CD138− subsets. Incubation of highly purified TCRβ+CD4+CD138− and TCRβ+CD8+CD138− for 5 days indicated that a substantial number of CD138+ cells derived from CD4+ cells. As shown in Figure 2E and paralleling the kinetics of CD138 expression among the TCRβ+CD138− population in Figure 2D, a significant increase in CD138 expression was observed among the CD4+ cells on day 1 which plateaued on day 3, while the increase in CD138+ expressing population among the CD8+ cells remained limited (Figure 2E). These results suggested that a portion of TCRβ+CD4+CD138− cells expresses CD138 after culture.

**Figure 2.**
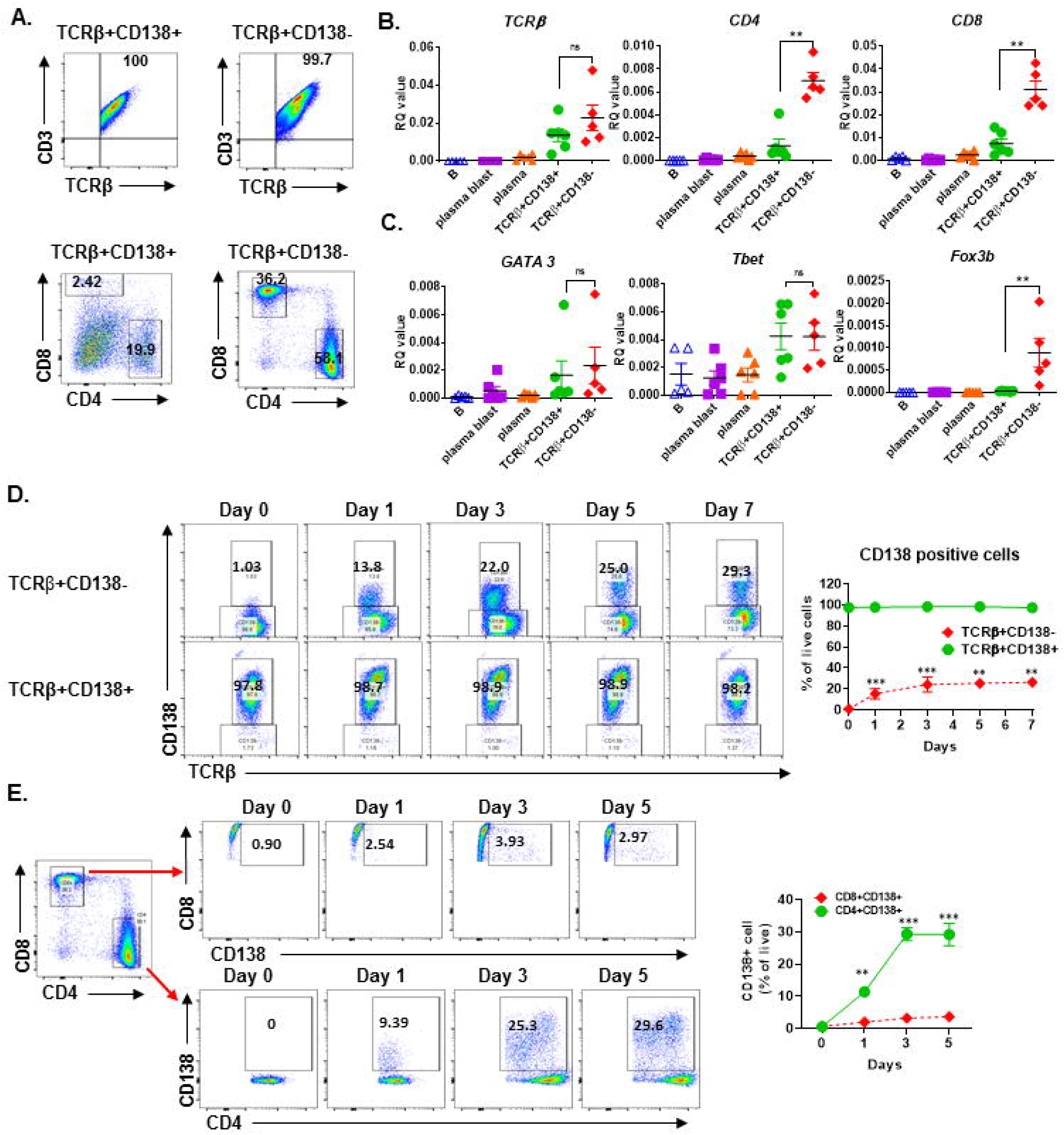
Most of the TCRβ+CD138+ cells are CD4 and CD8 negative while some derive from CD4+ T cells. **(A)** Splenocytes were harvested from 14 weeks old MRL/Lpr mice. After pre-gating live and single cells, TCRβ+CD138+ and TCRβ+138− cells were further analyzed for CD3, CD4 and CD8 expression by flow cytometry. Representative pseudocolor plots from out of 5 mice are shown. **(B and C)** Splenocytes were collected from 14 weeks old MRL/Lpr mice. CD19+ B cells, CD19+CD138− plasmablasts, CD19-CD138+ plasma cells, CD19-TCRβ+CD138− cells, and CD19-TCRβ+CD138+ cells were sorted by flow cytometry. The mRNA expression levels of cell surface molecules *TCR*β, *CD4*, *CD8* **(B)** and transcription factors *GATA3*, *Tbet*, and *Foxp3* **(C)** were quantified in Q-PCR. Mean ± SD of 5 to 6 mice from three independent experiments are plotted. ns, not significant. *p<0.05, **p<0.01. **(D)** Splenocytes were collected from 14 weeks old MRL/Lpr mice. After removing CD19+ B cells, TCRβ+CD138− and TCRβ+CD138+ cells were sorted with magnetic beads and then cultured separately for 5 days. Frequencies of CD138− expressing TCRβ+ cells were quantified by flow cytometry. Representative pseudocolor plots are shown. Mean percentages ± SD of 6 mice from three independent experiments are plotted. **p<0.01, and ***p<0.001 between TCRβ+CD138− and TCRβ+CD138+ cells. **(E)** Splenocytes were collected from 14 weeks old MRL/Lpr mice. After removing CD19+ B cells, TCRβ+CD138-CD4+ and TCRβ+CD138-CD8+ cells were sorted with magnetic beads and then cultured separately for 5 days. Frequencies of CD138 expressing CD4+ and CD8+ T cells were measured by flow cytometry. Mean percentages ± SD of 6 mice from three separate experiments are plotted. **p<0.01, and ***p<0.001 for TCRβ+CD138− vs TCRβ+CD138+ cells.

### TCRβ+CD138+ cells respond less to TCR engagement and PMA/ionomycin activation compared to CD138 counterparts

After TCR activation, T cells undergo a series of events, including up-regulation of cell surface markers, proliferation, apoptosis, and cytokine secretion [29]. We assessed the differences in proliferation of TCRβ+CD138+ and TCRβ+CD138− cells following stimulation with anti-CD3/CD28 antibodies. Compared to TCRβ+CD138− cells, TCRβ+CD138+ cells proliferated significantly less at 48− and 72-hour time points (Figure 3A). Stimulated cells were also assessed for apoptosis by flow cytometry. Compared to TCRβ+CD138− cells, TCRβ+CD138+ cells had higher number of live cells accompanied by lower number of early apoptotic and necrotic cells (Figure 3B). The early and late activation state of TCR-stimulated T cells were assessed by CD69 and CD25 expression, respectively [30]. Overall, anti-CD3/CD28 antibody-induced activation of TCRβ+CD138+ cells were delayed compared to TCRβ+CD138− cells. At 24-hour time point, CD69 expression was higher on TCRβ+CD138− cells than on TCRβ+CD138+ cells, but by 48 hours its expression decreased compared to TCRβ+CD138+ cells (Figure 3C). Conversely, the increase in CD69 expression on TCRβ+CD138+ cells peaked with a delay at 48 hours. A similar delay in the activation markers were observed in PMA/ionomycin-stimulated TCRβ+CD138+ cells (Supplemental Figure 3A). In SLE, activated T cells participate in the inflammatory process through the production of cytokines such as IFNγ, TNFα and IL-17 [2–4, 7, 31, 32]. We found that after TCR engagement and PMA/ionomycin stimulation, more than 90% of TCRβ+CD138− cells were positive for IFNγ, while less than 30% of TCRβ+CD138+ cells produced IFNγ (Figure 3D, Supplemental Figure 3B and C). Similarly, TCRβ+CD138+ cells expressed less TNFα compared to TCRβ+CD138− cells (Supplemental Figure 3B). Unlike IFNγ and TNFα, IL-17 production was not different between the two subsets after TCR stimulation. However, TCRβ+CD138+ cells produced less IL-17 than TCRβ+CD138− cells after PMA/ionomycin stimulation (Figure 3D, Supplemental Figure 3B and C). These results indicate that the phenotype of TCRβ+CD138+ cells activated with either PMA/ionomycin or through TCR are markedly different than their CD138 deficient counterparts.

**Figure 3.**
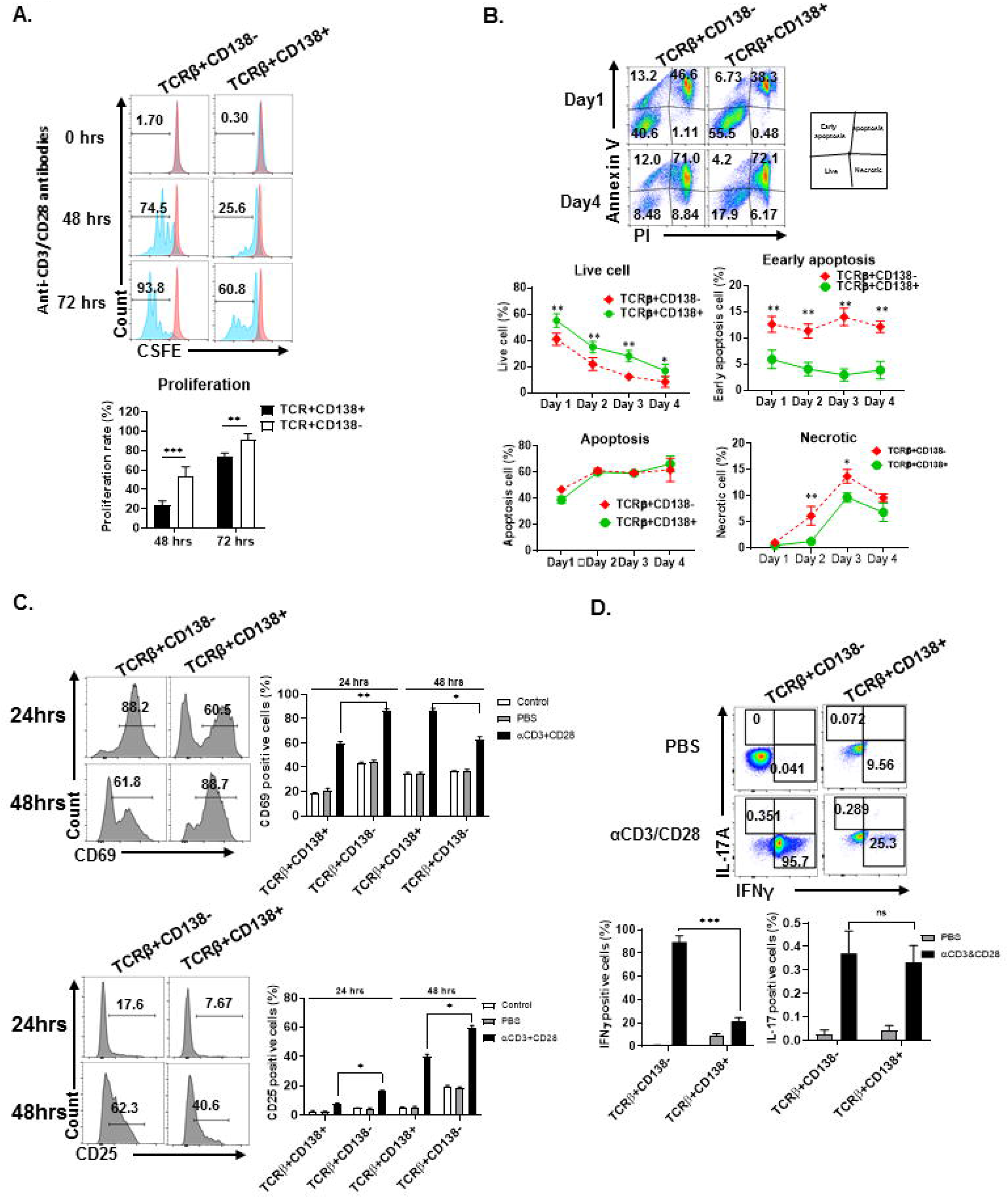
TCRβ+CD138+ cells proliferate less and resist early apoptosis after activation. **(A**) Splenic TCRβ+CD138+ and TCRβ+CD138− were sorted with magnetic beads and pre-stained with CSFE prior to stimulation with anti-CD3/CD28 antibodies for 3 days. The proliferation was assessed by flow cytometry. Representative histogram images are shown. Mean percentages ± SD of 7 mice from three separate experiments are plotted. **p<0.01, ***p<0.001. **(B)** Sorted splenic TCRβ+CD138+ and TCRβ+CD138− were stimulated for 4 days with anti-CD3/CD28 antibodies. Live, early apoptotic, apoptotic and necrotic cells were measured by flow cytometry. Representative pseudocolor plots are shown. Mean percentages ± SD of 6 mice from three independent experiments are plotted. *p<0.05, and **p<0.01 for TCRβ+CD138− vs TCRβ+CD138+ cells. **(C)** Sorted splenic TCRβ+CD138+ and TCRβ+CD138− cells were activated with anti-CD3/CD28 antibodies for 24 or 48 hours. Cells were stained with CD69 and CD25 antibodies to assess activation kinetics in flow cytometry. Representative histograms images indicating the frequency of positive cells are shown. Mean percentages ± SD of 7 mice from three independent experiments are plotted. *p<0.05, **p<0.01. **(D)** Sorted splenic TCRβ+CD138+ and TCRβ+CD138− cells were activated with anti-CD3/CD28 antibodies for 16 hours. Representative pseudocolor plots show intracellular IFN-γ, and IL-17 staining. For each cytokine, mean ± SD of 5 mice are plotted. ns, not significant. ***p<0.001.

### TCR-stimulated TCRβ+CD138+ cells are less efficient in activating B cells than TCRβ+CD138− cells

An important function of T cells is to enhance antibody-mediated immunity by driving B cell proliferation and development into long lived memory B cells or antibody-secreting plasma cells [33]. To investigate the capacity of TCRβ+CD138+ to help B cells, we co-incubated splenic B cells purified from 6 weeks old (disease free) MRL/Lpr mice with splenic TCRβ+CD138+ or TCRβ+CD138− cells, which were isolated from 10 to 12 weeks old MRL/Lpr mice in the presence of anti-CD3/CD28 antibodies and determined the proliferation of B cells (Supplemental Figure 4A). We found significantly less proliferation of B cells co-cultured with TCRβ+CD138+ cells than those co-cultured with TCRβ+CD138− cells (Figure 4A). Moreover, TCRβ+CD138+ cells were less efficient in inducing the generation of plasma cells (Figure 4B) and the production of IgM and IgG than TCRβ+CD138− cells (Figure 4C).

**Figure 4.**
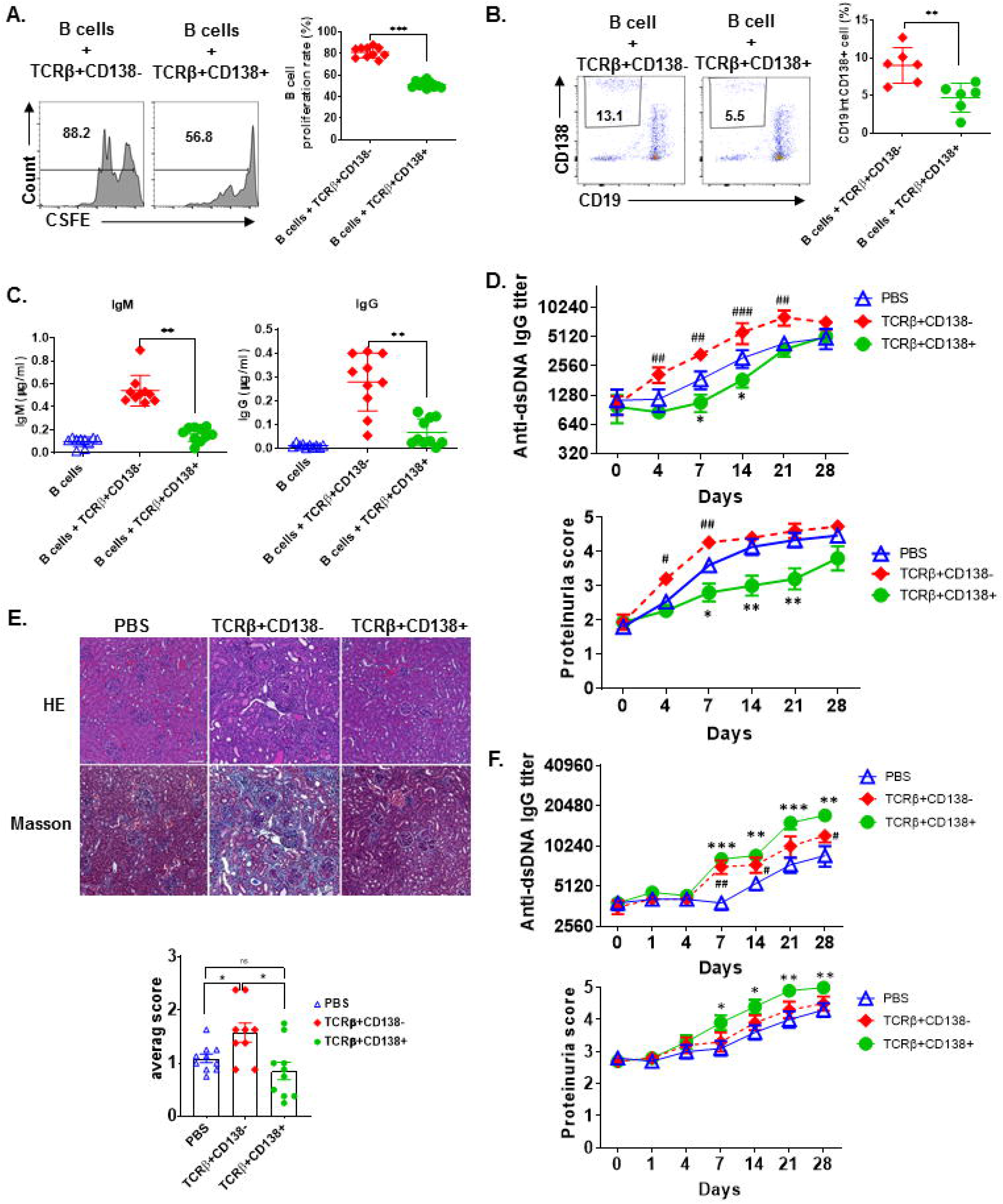
TCRβ+CD138+ cells are unable to promote lupus diseases development in young MRL/Lpr mice. **(A to C)** Sorted splenic TCRβ+CD138+ and TCRβ+CD138− cells from 10 to 12 weeks old MRL/Lpr mice were co-cultured with purified splenic B cells from 6 weeks old mice in the presence of anti-CD3/CD28 antibodies for 5 days. After gating-out TCRβ+ T cells, the proliferation of B cells **(A)** and the frequency of plasma cells (CD19^int^CD138+) **(B)** were measured by flow cytometry. Mean ± SD of 10 mice (A) or 6 mice (B) from three independent experiments are plotted. **p<0.01, ***p<0.001. **(C)** Culture supernatants from the above co-culture experiment were analyzed for total IgM and IgG concentrations by ELISA. Mean ± SD of 10 from mice from three independent experiments are plotted. **p<0.01. **(D and E)** Splenic TCRβ+CD138+ and TCRβ+CD138− cells were sorted from 10 to 12 weeks old MRL/Lpr mice and then adoptively transferred into 7 to 8 weeks old MRL/Lpr mice without disease symptoms. **(D)** Autoreactive IgG antibody (dsDNA) and proteinuria levels were measured on indicated days. Mean ± SEM of 15 mice from three independent experiments are plotted. #p<0.05, ##p<0.01, ###p<0.001, PBS vs TCRβ+CD138− cells; *p<0.05, **p<0.01, PBS vs TCRβ+CD138+ cells. **(E)** Kidneys were collected two weeks after the transfer of cells and histopathological evaluations were performed on H&E and Masson stained specimens. Average pathology scores of 10 mice from two separate experiments are plotted. ns, not significant, *p<0.05. **(F)** Splenic TCRβ+CD138+ and TCRβ+CD138− cells were sorted from 10 to 12 weeks old MRL/Lpr mice and then adoptively transferred into 11 to 12 weeks old MRL/Lpr mice with existing disease symptoms. Autoreactive IgG antibody (dsDNA) and proteinuria levels were measured on indicated days. Mean ± SEM of 10 mice from two independent experiments are plotted. #p<0.05, ##p<0.05, PBS vs TCRβ+CD138− cells; *p<0.05, **p<0.01, ***p<0.001, PBS vs TCRβ+CD138+ cells.

To further characterize TCRβ+CD138+ cells, we tested the immunomodulatory effect of TCRβ+CD138+ cells in SLE by adoptively transferring TCRβ+CD138+ or TCRβ+CD138− cells into MRL/Lpr mice and evaluating the disease progression. We conducted the adoptive transfer experiments in two different recipient mice age groups, 7 to 8 weeks old mice with minimal lupus symptoms and 11 to 12 weeks old mice with established lupus symptoms. We chose these two age groups to assess whether the CD138-expressing cells may impact the lupus progression differently in MRL/Lpr mice from different stages of disease. To our surprise, TCRβ+CD138+ cells had completely opposite effect in the recipient mice at different ages. The 7 to 8 weeks old recipient mice without lupus symptoms manifested slower progression of disease when they were transferred with TCRβ+CD138+ cells from 10 to 12 weeks old MRL/Lpr (sick) mice, compared to those that were injected with PBS. The increase in anti-dsDNA antibody and proteinuria levels were slower compared to those that were injected with PBS (Figure 4D, Supplemental Figure 4B). Kidney histopathological findings showed the most severe changes in recipients of TCRβ+CD138− group, including end stage glomeruloscrerosis with severe inflammation and interstitial fibrosis. The glomerular histopathological changes of TCRβ+CD138+ cell-recipient mice were reduced as compared to PBS-injected mice two weeks after the transfer of cells, but the difference in histopathological scores did not reach statistical significance (Figure 4E). However, histopathological changes were significantly less in TCRβ+CD138+ cell-injected mice than those injected with TCRβ+CD138− cells. In sharp contrast to its effect in young recipient mice, TCRβ+CD138+ cells significantly increased the development of anti-dsDNA antibodies and proteinuria in older MRL/Lpr mice with existing disease symptoms (Figures 4F, Supplemental Figure 4C). The TCRβ+CD138− cells accelerated the disease progression in recipient animals from both age groups but their impact on older recipient mice was not as pronounced as those that received TCRβ+CD138+ cells (Figure 4D to F, Supplemental Figure 4B and C). Together with the *in vitro* B cell co-incubation data (Figure 4A to C), the adoptive transfer experiments in young MRL/Lpr mice suggest an immunosuppressive function for TCRβ+CD138+ cells on B cell differentiation. However, TCRβ+CD138+ cells contribute to the acceleration of disease progression if the recipient host has established lupus disease.

### TCRβ+CD138+ cells augment autoreactive B cell responses when auto-antigens are present

Next, we sought to explore the underlying mechanism for the dramatic difference in the disease-stage-specific impact of TCRβ+CD138+ cells to lupus progression in the adoptive transfer experiments. An important distinction between the young and older recipient MRL/Lpr mice in the adoptive transfer experiments is the exposure of the immune system of older mice (11 to 12 weeks old), but not the younger (7 to 8 weeks old) mice to self-antigens as a result of apoptosis [34]. As observed with the younger mice in the adoptive transfer experiments, *in vitro* TCR stimulation of TCRβ+CD138+ cells also resulted in less activation of autoreactive B cells than under culture conditions containing TCR-stimulated TCRβ+CD138− cells. A common feature of these *in vitro* and *in vivo* experiments is the absence of auto-antigens in the system. Earlier studies has highlighted the importance of B cells in presenting auto-antigens to T cells in activating autoreactive T cells [35]. We therefore repeated the *in vitro* co-culture experiments in the presence of apoptotic DNA instead of anti-CD3/CD28 antibodies. Interestingly, when B cells from 12 weeks old MRL/Lpr mice with established disease were co-incubated with TCRβ+CD138+ cells from age matched MRL/Lpr mice in the presence of DNA from apoptotic cells, culture supernatants contained significantly higher autoreactive (Figure 5A) and total IgG and IgM antibodies than those co-cultured with TCRβ+CD138− cells (Supplemental Figure 5A). Consistent with the difference in antibody responses, higher percentage of plasma cells (CD3-CD19-CD138+) emerged from the cultures containing TCRβ+CD138+ cells than from the cultures with TCRβ+CD138− cells (Figure 5B). Increased plasma cell generation and autoreactive antibody production with TCRβ+CD138+ cells were not restricted to self-DNA-containing cultures, because replacement of self-DNA with the auto-antigen SM also resulted in higher percentage of plasma cell development and increased production of anti-SM antibodies with TCRβ+CD138+ cells than with TCRβ+CD138− cells (Figure 5C, Supplemental Figure 5B). Thus, regardless of the nature of the autoantigen, TCRβ+CD138+ cells are more potent in aiding autoreactive B cells to produce self-reactive antibodies than TCRβ+CD138− cells.

**Figure 5.**
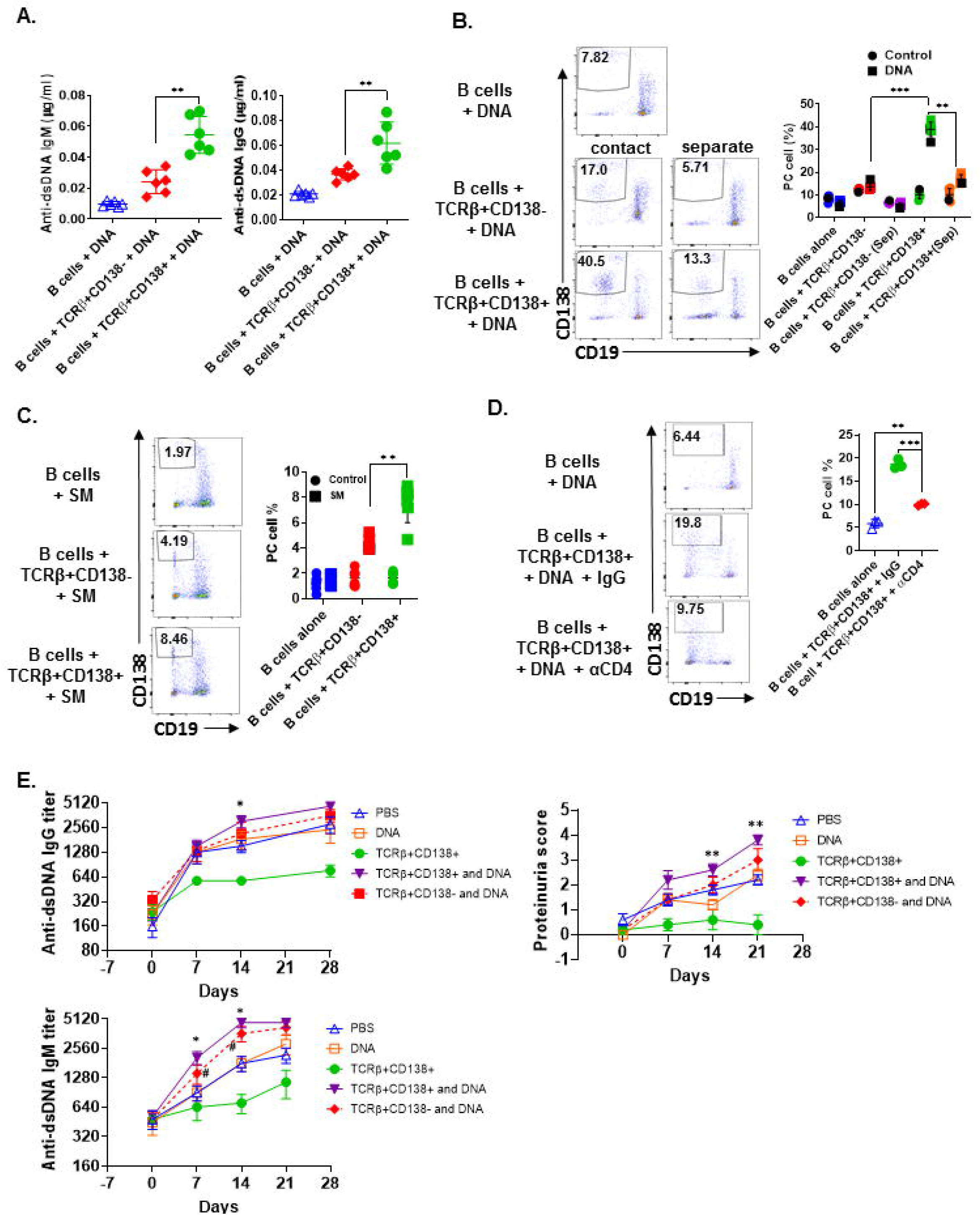
TCR+CD138+ cells activate autoreactive B cells when auto-antigens are included in the culture. **(A-D)** Sorted splenic TCRβ+CD138+ and TCRβ+CD138− cells from 12 weeks old MRL/Lpr mice were co-cultured with purified splenic B cells from the same mice for 5 days. **(A)** DNA was included in the co-cultured cells and culture supernatant anti-dsDNA IgM and IgG antibodies were measured by ELISA. Mean ± SD of 6 mice from three independent experiments are plotted. **p<0.01. **(B)** T and B cells were incubated as mixed or separated with Transwell® (Sep) in the presence of DNA. After gating-out TCRβ+ T cells, the differentiation of B cells into plasma cells (CD19^int^CD138+) was quantified by flow cytometry. Mean percentages ± SD of 6 mice from three independent experiments are plotted. **p<0.01, ***p<0.001. **(C)** SM was included in the co-cultured cells and differentiation of B cells into plasma cells (CD19^int^CD138+) were quantified by flow cytometry after gating-out TCRβ+ T cells. Mean percentages ± SD of 6 mice from three experiments are plotted. **p<0.01. **(D)** Cells were incubated in the presence of antibody against CD4 or control IgG and the differentiation of B cells into plasma cells was quantified by flow cytometry. Mean percentages ± SD of three independent experiments are plotted. **p<0.01, ***p<0.001. **(E)** Splenic TCRβ+CD138+ and TCRβ+CD138− cells from 10 to 12 weeks old MRL/Lpr mice were sorted and then adoptively transferred into 5 to 6 weeks old MRL/Lpr mice with or without DNA. Mice that received PBS or DNA only served as control. Serum anti-dsDNA IgG and IgM antibody as well as proteinuria levels at indicated days were measured. Mean ± SEM of 5 mice are plotted. #p<0.05, DNA vs TCRβ+CD138− and DNA; *p<0.05, **p<0.01, DNA vs TCRβ+CD138+ and DNA.

We next sought to determine whether TCRβ+CD138+ cell-enhanced B cell differentiation is mediated by direct cell contact, especially because the majority of the MRL/Lpr mice TCRβ+CD138+ cells were CD4 and CD8 negative (Figure 2A). Separation of B and TCRβ+CD138+ cells in the co-culture experiments with a Transwell® system resulted in a significant reduction in plasma cell generation compared to cells cultured without a Transwell® system (Figure 5B). Moreover, although CD4+ cells constituted only approximately 20% of the TCRβ+CD138+ cells, the B cell help provided by TCRβ+CD138+ cells required CD4 because inclusion of anti-CD4 blocking antibodies in the co-culture system severely reduced plasma cell development and production of anti-dsDNA IgG and IgM antibodies (Figure 5D, Supplemental Figure 5C). These experiments established the TCRβ+CD4+CD138+ cells as more potent autoreactive B cell-activating T cell subset when self-antigens are present in the culture environment.

### TCRβ+CD138+ cells promote disease in MRL/Lpr mice only when self-antigens are exposed

We showed that in order for TCRβ+CD138+ cells to augment autoreactive B cell responses, they need to be stimulated by B cell-presented self-antigens (Figure 5A to D). The fact that the recipient MRL/Lpr mice used in the adoptive transfer experiments in figure 4D were too young to have sufficient amounts of circulating self-antigens, such as DNA, could be the reason why the transferred TCRβ+CD138+ cells were not activated and did not exacerbate SLE symptoms in the recipient mice. Conversely, the acceleration of disease progression in older MRL/Lpr mice after the transfer of TCRβ+CD138+ cells could be due to the presence of circulating self-antigens. To test this possibility, we co-administered TCRβ+CD138+ cells with DNA into young (5 to 6 weeks old) MRL/Lpr mice. As observed previously (Figure 4D), the increase in anti-dsDNA IgG and IgM antibodies as well as proteinuria were significantly slower in mice injected only with TCRβ+CD138+ cells compared to PBS-injected mice or control mice injected with DNA only (Figure 5E). In contrast, and as hypothesized, anti-dsDNA antibody and proteinuria levels were significantly higher in mice co-administered with DNA and TCRβ+CD138+ cells than in control mice injected with DNA only (Figure 5E). Taken together, TCRβ+CD138+ cells can modulate lupus development in MRL/Lpr mice in a disease-stage-dependent manner; they slow down the symptoms prior to the emergence of self-antigens and accelerate the disease progression when self-antigens are exposed.

### TCR+CD138+ cells are central memory biased T cells

Abnormal accumulation and differentiation of memory T cells have been reported in lupus patients [13]. Memory T cells confer immediate protection and mount recall responses upon reencounter with antigens. Since adoptively transferred TCRβ+CD138+ cells promoted disease progression in an autoantigen dependent manner (Figure 5), we asked whether TCRβ+CD138+ cells have a memory T cell phenotype. The circulating memory T cell compartments are divided into Tcm and effector memory T cells (Tem) subsets based on the expression of cell surface molecules, such as CD44, CD62L and CCR7 [36]. We first characterized the expression of CD44 and CD62L on splenic TCRβ+CD138+ and TCRβ+CD138− cells of 12 weeks old MRL/Lpr mice. Using these two markers, we identified CD44-CD60L+ naïve T (Tn) cells, CD44+CD60L- Tem and CD44+CD60L+ Tcm subsets. Among the TCRβ+CD138− cells, appo 10% were Tn, 50% Tem and 40% Tcm (Figure 6A, Supplemental Figure 6A). Interestingly, Tcm cells comprised the vast majority (95%) of the population among the TCRβ+CD138+ cells (Figure 6A, Supplemental Figure 6A). Typically, memory T cells are either CD4+ or CD8+ [37], but in MRL/Lpr mice, the majority of the CD138-expressing CD4+, CD8+ as well as CD4-CD8− cells exhibited Tcm memory phenotype, based on the elevated expression of CD44 and CD60L (Supplemental Figure 6B).

**Figure 6.**
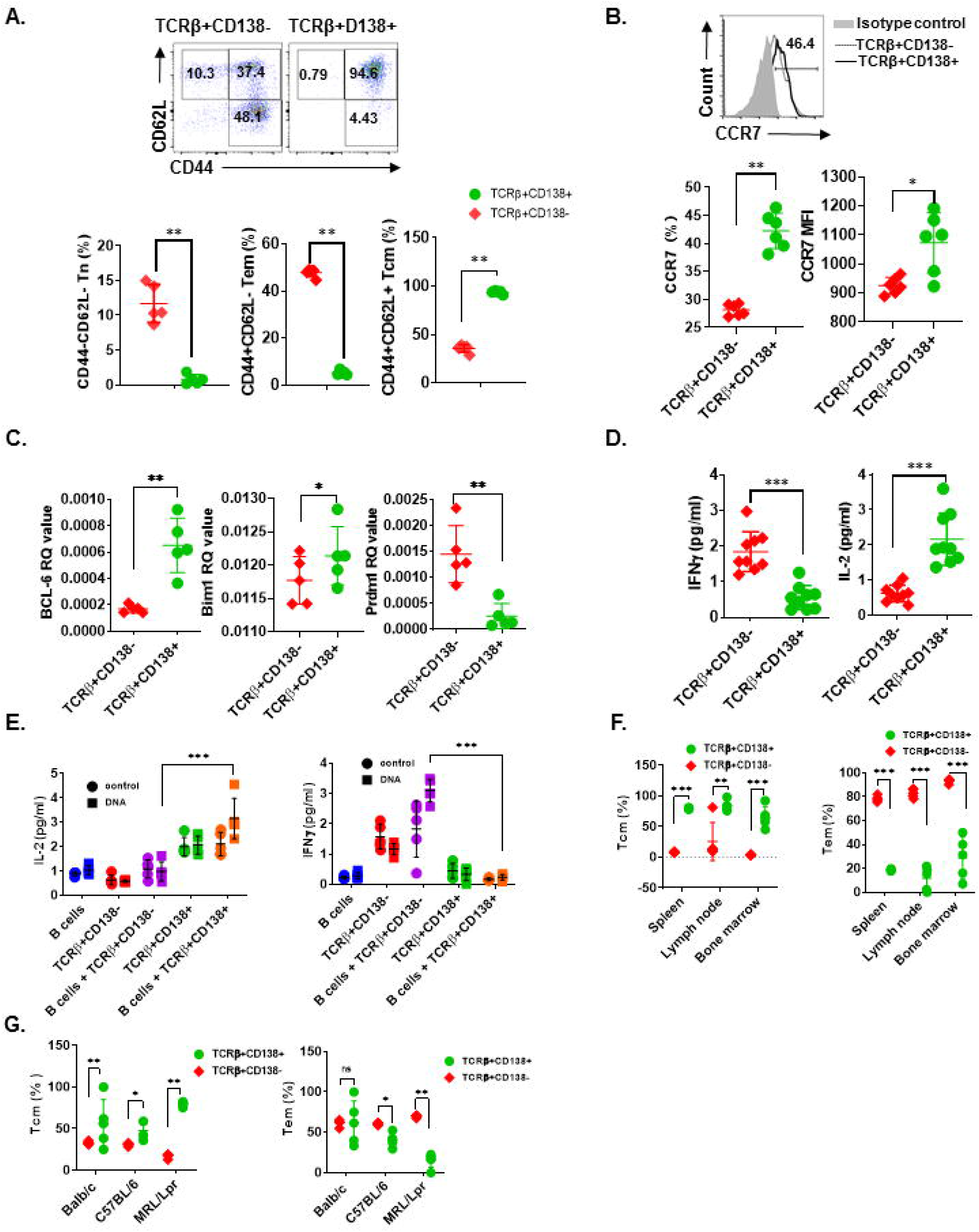
CD138+ T cells exhibit central memory T cell phenotype. **(A and B)** Splenocytes were collected from 12 weeks old MRL/Lpr mice. **(A)** The expression of CD44 and CD62L on TCRβ+CD138− and TCRβ+CD138+ cells were measured by flow cytometry. Representative pseudocolor plots are shown. Mean percentages ± SD of 5 mice from two separate experiments are plotted. **p<0.05. **(B)** Representative flow cytometry histogram of CCR7 expression on TCRβ+CD138− and TCRβ+CD138+ cells are shown. Mean ± SD percentages and MFIs for CCR7 expression from 6 mice in two separate experiments are plotted. *p<0.05, **p<0.01. **(C)** Splenic TCRβ+CD138+ and TCRβ+CD138− cells were sorted from 12 weeks old MRL/Lpr mice and *BCL-6, Bim1* and *Prdm1* mRNA were quantified by Q-PCR. Mean ± SD values of 5 mice from two separate experiments are plotted. *p<0.05, **p<0.01. **(D)** Splenic TCRβ+CD138+ and TCRβ+CD138− cells were sorted from 12 weeks old MRL/Lpr mice and cultured with anti- CD3/CD28 antibodies overnight. Culture supernatant IFNγ and IL-2 levels were measured by ELISA. Mean ± SD of 9 mice from three independent experiments are plotted. ***p<0.001. **(E)** Splenic TCRβ+CD138+, TCRβ+CD138− cells were sorted from 12 weeks old MRL/Lpr mice and co-cultured with sorted B cells from the same mice in the presence of DNA for 3 days. Culture supernatant IFNγ and IL-2 levels were measured by ELISA. Mean ± SD of 5 mice from two separate experiments are plotted. ***p<0.001. **(F)** Cells were collected from spleen, lymph nodes and bone marrows of 12 weeks old MRL/Lpr mice. Frequencies of Tcm (CD44+CD60L+) and Tem (CD44+CD60L−) cells among TCRβ+CD138− and TCRβ+CD138+ cells were measured in flow cytometry. Mean percentages ± SD of 5 mice from two independent experiments are plotted. **p<0.01, ***p<0.001. **(G)** Splenocytes were collected from 12 weeks old Balb/c, C57BL/6 and MRL/Lpr mice. Frequencies of Tcm (CD44+CD60L+) and Tem (CD44+CD60L−) cells among TCRβ+CD138− and TCRβ+CD138+ cells were measured in flow cytometry. Mean percentages ± SD of 5 mice from two independent experiments are plotted. ns, not significant, *p<0.05, **p<0.01.

The chemokine receptor CCR7, which enables cells to home to secondary lymphoid organs where they encounter antigen, are highly expressed on Tcm cells [38]. Further confirming their Tcm phenotype, TCR+CD138+ cells expressed higher CCR7 levels than TCRβ+CD138− cells (Figure 6B). The Tcm and Tem memory subsets can also be distinguished based on the expression of transcription factors Bcl-6 and Bim for Tcm and Prdm1 for Tem, respectively [39, 40]. We found higher *Bcl-6* and *Bim1* but lower *Prdm1* mRNA expression in TCRβ+CD138+ cells, compared to TCRβ+CD138− cells (Figure 6C). We also assessed Tcm phenotype based on IFNγ and IL-2 production because in humans, Tcm cells produce more IL-2, while Tem cells are distinguished by high IFNγ and TNFα production [41]. Consistent with data in humans upon stimulation with anti-CD3/CD28 antibodies, MRL/Lpr mice TCRβ+CD138+ cells secreted significantly higher IL-2 but lower IFNγ than TCRβ+CD138− cells (Figure 6D). The Tcm specific IL-2 and IFNγ production profile in TCRβ+CD138+ cells were also observed in co-culture system with autoantigens and B cells (Figure 6E). Moreover, we found that in MRL/Lpr mice, the Tcm phenotype of TCRβ+CD138+ cells were not restricted to the spleen or the age because cells isolated from various organs from mice at different ages, all exhibited the Tcm phenotype (Figure 6F, Supplemental Figure 6B and C). Collectively, these data established the memory phenotype of TCRβ+CD138+ cells as Tcm in MRL/Lpr mice. Finally, to assess whether the Tcm phenotype of TCRβ+CD138+ is unique to MRL/Lpr mouse, we compared the memory phenotype of TCRβ+CD138+ cells from Balb/c and C57BL/6 mice to those of MRL/Lpr mice. Again, regardless of the mouse strain, TCRβ+CD138+ were CD44+CD62L+ Tcm cells (Figure 6G). Thus, the Tcm phenotype of TCR+CD138+ cells is conserved among different mouse strains.

## DISCUSSION

In this study, we uncovered a disease-dependent accumulation of TCRβ+CD138+ cells in various organs of lupus-prone MRL/Lpr mice, which are overwhelmingly positive for B220, but negative for CD4 and CD8 expression. Although these cells are less efficient in responding to non-specific T cell stimuli, such as PMA/ionomycin and anti-CD3/CD28 antibody engagement, they are more potent in aiding antibody production from autoreactive B cells *in vitro* as well as *in vivo* when autoantigens were present. Further characterization established that these cells have a Tcm phenotype based on high expressiong levels of CD62L, CD44, CCR7 and Bcl-6.

CD138 is widely expressed on epithelial cells as well as on other adherent cells, but its expression on normal lymphoid cells has been thought to be restricted to plasma cells and pre-B cells. However, recent studies reported that CD138 is also present on NKT17 and GC B cells, where it may be involved in host defense or autoimmunity through IL-17 secretion or binding to death receptor 6 on Tfh cells [22, 42]. In aged C3H mice, accumulation of CD138 expressing T cells was shown to be restricted to the gut epithelium, although they can expand to peripheral organs, such as lymph nodes and spleen when Fas ligand (gld) is ablated. Similar to C3H gld mice, a large population of CD138-expressing T cells accumulate in peripheral organs of Fas receptor mutant μMT/Lpr, B6/Lpr [25] and MRL/lpr mice (Figure 1A). As previously described, the majority of CD138+ T cells also express CD3 and B220 and are negative for CD4 and CD8 (dnT). Previous reports suggested that dnT cells derive from exhausted autoreactive CD8+ cells or continuously stimulated CD8+ cells [43, 44]. Differing from these reports, our observations reveal that a substantial portion of the TCRβ+CD138+ cells in MRL/Lpr mice are converted from CD4+ cells rather than CD8+ cells, as *in vitro* cultured CD4+ cells, but not CD8+ cells, resulted in the accumulation of CD138+ cells (Figure 2D).

Studies in normal mice or autoimmune-prone lpr mice have shown that dnT cells are able to dampen CD4+ and CD8+ T cell-mediated autoimmune responses both *in vitro* and *in vivo* [45, 46]. Consistent with these studies, adoptively transferred TCRβ+CD138+ T cells, of which the majority were CD4-CD8− cells, slow down the disease progression in young MRL/Lpr mice when auto-antigens are not exposed. The TCRβ+CD138+ cell-mediated-suppression of disease progression is unlikely to be due to the apoptosis of host CD4+ and CD8+ T cells induced by the transferred cells as this mechanism requires functional Fas-FasL interaction [46]. Suppression of dnT cell-mediated alloimmune responses are also attributed to elevated levels of perforin and granzyme B produced by these cells [47]. This mechanism is also unlikely to be at play in the slowing down of disease progression by TCRβ+CD138+ cells in young MRL/Lpr mice because comparable levels of perforin and granzyme B production are detected in TCRβ+CD138+ and TCRβ+CD138− cells (Supplemental Figure 7). Although the exact molecular mechanism and cascade of events need to be deciphered , we augur that the suboptimal proliferation capacity of TCRβ+CD138+ cells as well as their diminished production of pathogenic cytokines, IL-17, TNFα and IFNγ [48] may be responsible for the delay in disease progression afforded by these cells in young MRL/Lpr mice.

In active lupus patients or NZBxSWR mice, CD4+ as well as dnT cells augment the production of anti-dsDNA antibodies when they are co-cultured with oligoclonal autoreactive B cells [35, 49, 50]. Consistent with these early observations, both TCRβ+CD138+ and TCRβ+CD138− cells enhance plasma cell development and amplify autoreactive antibody production from MRL/Lpr mice B cells when auto-antigens were present. However, TCRβ+CD138+ were more potent than TCRβ+CD138− cells in activating autoreactive B cells both *in vitro* and *in vivo*. Interestingly, CD4-expressing TCRβ+CD138+ cells are responsible for the activation of autoreactive B cells, despite comprising less than 20% percent of the total TCRβ+CD138+ population. This rapid recall response upon antigen re-encounter is a typical characteristic of memory T cells [36, 41]. Indeed, compared to TCRβ+CD138− cells, over 90% of TCRβ+CD138+ cells are CD44+CD62L+ Tcm cells. Although the other Tcm marker CCR7 is also higher on TCRβ+CD138+ cells than on TCRβ+CD138− cells, the frequency of CCR7-expressing TCRβ+CD138+ cells is less than 50%. The discrepancy between the percentage of CD44+CD62L+ cells and CCR7+ cells may be due to possible loss of CCR7 expression on TCRβ+CD138+ cells after repeated exposure to autoantigens, a phenomenon reported for CCR7+CD27+ memory T cells in lupus patients which loose CCR7 expression following repeated stimulation [13]. Of note, the Tcm characteristic of TCRβ+CD138+ cells is independent of disease activity as TCRβ+CD138+ cells from healthy MRL/Lpr mice as well as different healthy mouse strains also exhibit similar phenotype. Thus, TCRβ+CD138+ cells manifest Tcm phenotype, regardless of mouse strain and disease status.

Taken together, we have identified and characterized the phenotype of TCRβ+CD138+ cells in MRL/Lpr mice and unveiled a pathogenic role for TCRβ+CD138+ cells in lupus disease. Both *in vivo* adoptive transfer experiments and *in vitro* co-culture experiments underpin the role of TCRβ+CD138+ cells in activating autoreactive B cells when autoantigens are exposed. Although how CD138 expression renders TCRβ+ T cells more pathogenic is still enigmatic, the discovery of a novel Tcm subset with enhanced pathogenic properties may be useful in assessing and monitoring disease severity.

