## Supplemental Figure text for "Disease stage-specific pathogenicity of CD138 (syndecan 1)-expressing T cells in systemic lupus erythematosus"

**SUPPLEMENTAL FIGURES**

**Supplemental Figure 1. MRL/Lpr mice splenocytes contain a population of TCR**β **and CD138 double positive cells. (A)** Gating strategy for TCRβ+CD138+ cells is shown. MRL/Lpr mice splenocytes were stained with CD19, TCRβ and CD138 antibodies. After gating on live and single cells, CD138 expressing T cells (CD19-TCRβ+) were analyzed in flow cytometry. **(B)** Q-PCR analysis of CD138 mRNA expression from flow cytometry sorted TCRβ+CD138+ and TCRβ+CD138- cells. Mean ± SD of 5 mice from three experiments are plotted. **p<0.01 **(C)** Average spleen weight and serum anti-SM IgG titers of PBS and pristane treated Balb/c and C57BL/6 mice are plotted. **(D)** Increased TCR+CD138+ cells in pristane-induced lupus disease in Balb/c and C57BL/6 mice. The frequencies of TCRβ+CD138+ cells in pristane-treated Balb/c and C57BL/6 mice were determined in flow cytometry. Representative pseudocolor plots are shown. In experiments C and D, mean ± SD of 5 mice are shown. **p<0.01, ***p<0.001.

**Supplemental Figure 2.** **TCRβ+CD138+ cells do not express canonical B cell markers. (A and B)** Splenocytes were harvested from 10 to 12 weeks old MRL/Lpr mice. After pre-gating live and single cells, B220 **(A)**, BAFFR, BCMA and TACI **(B)** expression were assessed by flow cytometry. Representative pseudocolor **(A)** and histogram **(B)** plots of 5 mice from three experiments are shown. **(C to E)** CD19+ B cells, CD19+CD138- plasmablasts, CD19-TCRβ-CD138+ plasma cells, CD19-TCRβ+CD138- cells, CD19-TCRβ+CD138+ cells were sorted from spleens of 10 to 12 weeks old MRL/Lpr mice by flow cytometry, and the expression of canonical B cells molecules *CD19*, *IGHM* (IgM), *Myc,* *TNFRSF13C* (BAFFR) and *TNFRSF13B* (TACI) as well as CD21, CD23, CD40, CD80 and CD86 were quantified by Q-PCR **(C)** and by flow cytometry **(D)**, respectively. Mean ± SD of three experiments from 5 to 6 mice are plotted. ns, not significant, **p<0.01, ***p<0.001. **(E)** The same cells were also analyzed for the expression of transcription factors *BCL-6, Pax5, PU1, Prdm1, Irf4*, and *Xbp1* by Q-PCR. Mean ± SD of 5 to 6 mice from three experiments are plotted. ns, not significant, *p<0.05, **p<0.01

**Supplemental Figure 3. Slower activation kinetics and diminished cytokine production in TCRβ+CD138+ cells.** Splenic TCRβ+CD138+ and TCRβ+CD138- cells from 10-12 weeks old MRL/Lpr mice were sorted with magnetic beads. **(A)** Sorted cells were stimulated with PMA/ionomycin for 24 or 48 hours and the activation kinetics was assessed by measuring CD69 and CD25 expression in flow cytometry. Representative histogram images indicating the frequencies of CD69 and CD25 expressing cells are shown. Mean percentages ± SD of 7 mice from three experiments are plotted. **(B)** Sorted cells were stimulated with anti-CD3/CD28 antibodies and mRNA for *IFNγ*, *TNFα* and *IL-17* were quantified by Q-PCR. Mean ± SD of 5 mice from two experiments are plotted. **(C)** Sorted cells were stimulated with PMA/ionomycin, and the production of IFNγ, and IL-17 were measured by flow cytometry. Representative pseudocolor plots show intracellular IFN-γ, and IL-17 staining. Mean percentages ± SD of 5 mice from two experiments are plotted. ns not significant**,** *p<0.05, **p<0.01, ***p<0.001.

**Supplemental Figure 4. TCRβ+CD138+ cells are unable to promote B cell proliferation and *in vivo* auto-antibody production when adoptively transferred into young MRL/Lpr mice. (A)** Gating strategy for B cell proliferation in B and T cell co-culture system. Sorted splenic TCRβ+CD138+ and TCRβ+CD138- cells from 10 to 12 weeks old MRL/Lpr mice were co-cultured with purified splenic B cells from 6 weeks old MRL/Lpr mice with no disease symptoms in the presence of anti-CD3/CD28 antibodies for 5 days. The proliferation of CSFE labelled B cells were measured by flow cytometry. **(B and C)** Splenic TCRβ+CD138+ and TCRβ+CD138- cells from 10 to 12 weeks old weeks old MRL/Lpr mice were sorted and adoptively transferred into either 7 to 8 weeks old MRL/Lpr mice with minimal lupus symptoms **(B)** or 11 to 12 weeks old MRL/Lpr mice with established disease symptoms **(C)**. Serum anti-dsDNA IgM antibody levels on indicated days were measured by ELISA. Mean ± SEM of 15 mice from three experiments are plotted. *p<0.05 and **p<0.01 for PBS vs TCRβ+CD138-, #p<0.05 and ##p<0.01 for TCRβ+CD138+ vs TCRβ+CD138- cells.

**Supplemental Figure 5. TCRβ+CD138+ cells promote auto-antibody production in B-cell co-culture system when autoantigens are present. (A and B)** Sorted splenic TCRβ+CD138+ and TCRβ+CD138- cells from 12 weeks old MRL/Lpr mice were co-cultured with B cells from the same mice in the presence of DNA **(A)** or SM **(B)** for 5 days. **(A)** The IgM and IgG concentrations in the culture media were measured by ELISA. Mean ± SD of 6 mice from three experiments are plotted. *p<0.05, **p<0.01 **(B)** Culture supernatant total and SM specific IgM and IgG antibodies were measured by ELISA. Mean ± SD of 9 mice from three experiments are plotted. ns, not significant, *p<0.05, **p<0.01 **(C)** Sorted splenic TCRβ+CD138+ and TCRβ+CD138- cells from 12 weeks old MRL/Lpr mice were co-cultured with B cells from the same mice in the presence of DNA and blocking antibodies against CD4. After 5 days, culture supernatant total and dsDNA specific IgM and IgG antibodies were measured by ELISA. Mean ± SD of 6 mice from three experiments are plotted. ns, not significant**,** *p<0.05, **p<0.01, ***p<0.001.

**Supplemental Figure 6. TCRβ+CD138+ cells exhibit central memory T cell-phenotype. (A)** Gating strategy for the analysis splenic Tcm and Tem cells. **(B)** Splenocytes were collected from 3, 12 and 30 weeks old MRL/Lpr mice. The expression of CD44 and CD62L on CD4, and CD8 single positive or double negative cells were measured by flow cytometry. Representative pseudocolor plots are shown. Mean percentages ± SD for 5 mice from two experiments are plotted. ns, not significant**,** *p<0.05, **p<0.01, ***p<0.001 **(C)** Splenocytes, lymph nodes, and bone marrow were collected from 30 weeks old MRL/Lpr mice. The expression of CD44 and CD62L on CD4+, dnT, and CD8+ subsets of TCRβ+CD138- and TCRβ+CD138+ cells were measured by flow cytometry. Representative pseudocolor plots are shown. Mean percentages ± SD for 5 mice from two experiments are plotted. ns not significant**,** *p<0.05, **p<0.01, ***p<0.001.

**Supplemental Figure 7. Activated TCRβ+CD138+ cells express comparable levels of perforin and Granzyme B.** Sorted TCRβ+CD138- and TCRβ+CD138+ cells from spleens of 10 to 12 weeks old MRL/Lpr mice were cultured with anti-CD3/CD28 antibodies overnight. The expression of perforin and granzyme B were quantified by Q-PCR. Mean ± SD of 5 mice from two experiments are plotted. ns not significant.
