## Supplemental Figures for "Disease stage-specific pathogenicity of CD138 (syndecan 1)-expressing T cells in systemic lupus erythematosus"

### Slide 1
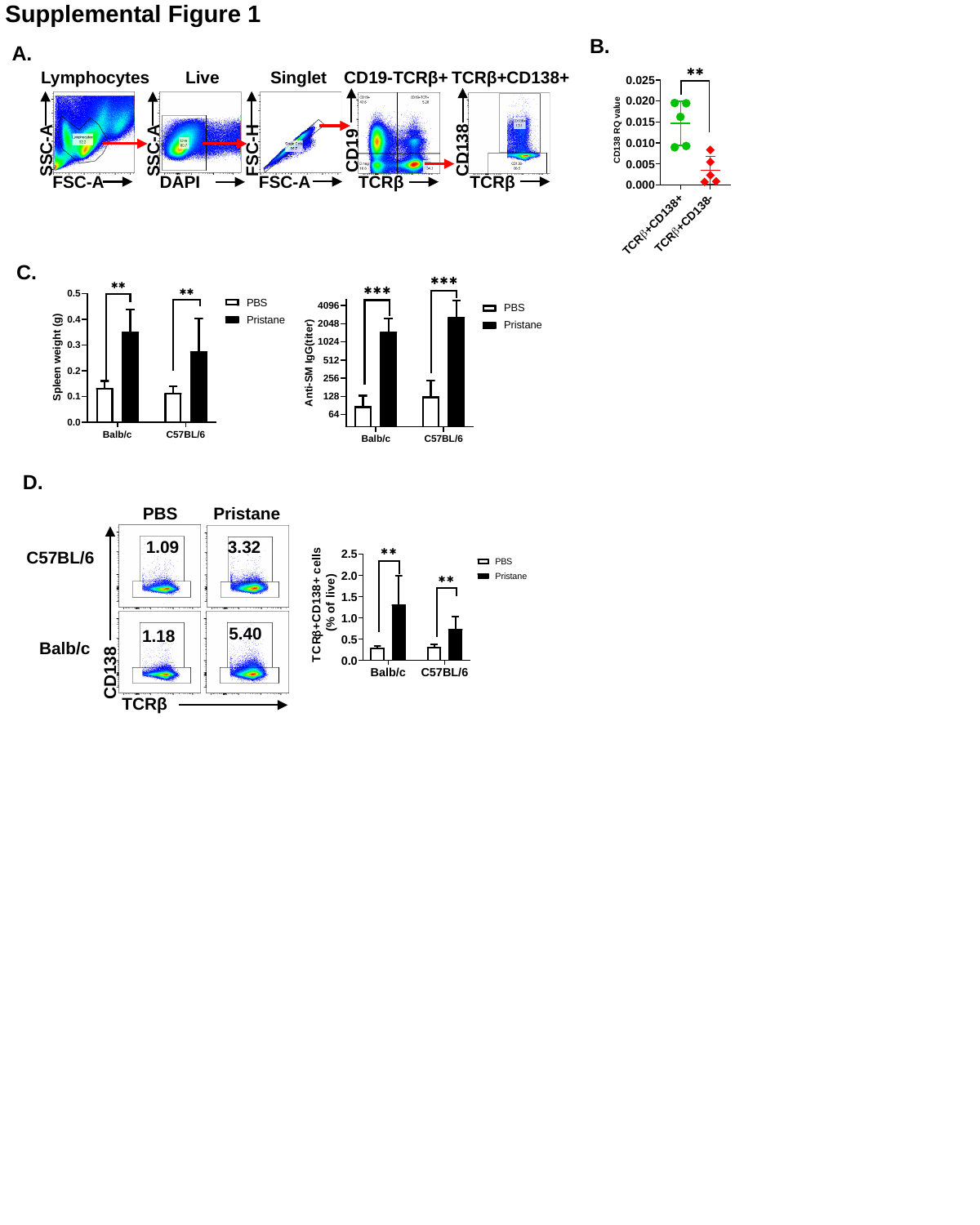

Supplemental Figure 1
B.
A.
Lymphocytes
Live
Singlet
CD19-TCRβ+
TCRβ+CD138+
FSC-H
CD138
SSC-A
CD19
SSC-A
FSC-A
TCRβ
FSC-A
TCRβ
DAPI
C.
D.
PBS
Pristane
1.09
3.32
C57BL/6
5.40
1.18
Balb/c
CD138
TCRβ

### Slide 2
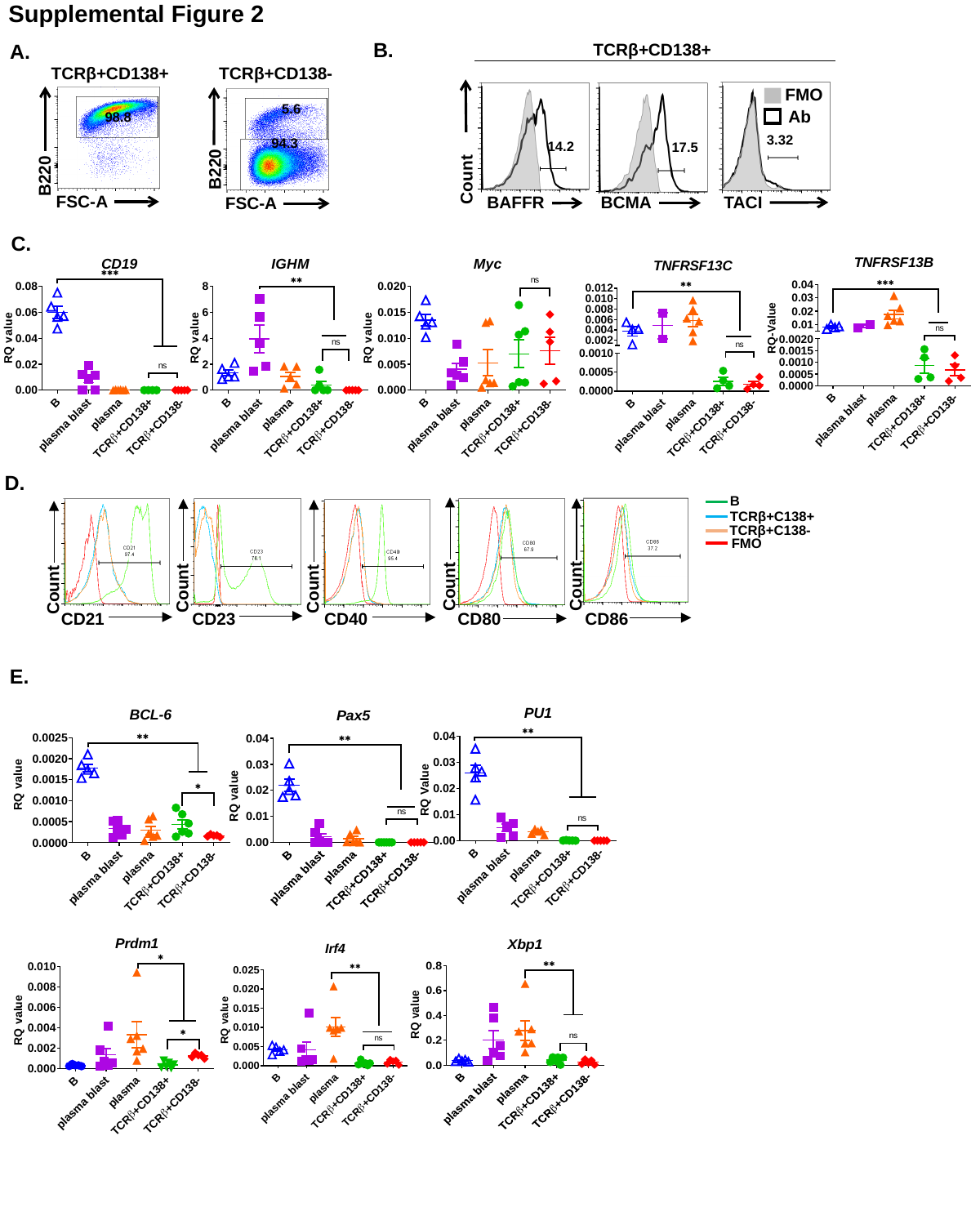

Supplemental Figure 2
B.
TCRβ+CD138+
A.
TCRβ+CD138+
TCRβ+CD138-
FMO
5.6
Ab
98.8
3.32
94.3
Count
14.2
17.5
B220
B220
FSC-A
BAFFR
BCMA
TACI
FSC-A
C.
D.
B
TCRβ+C138+
TCRβ+C138-
FMO
Count
Count
Count
Count
Count
CD21
CD23
CD40
CD80
CD86
E.

### Slide 3
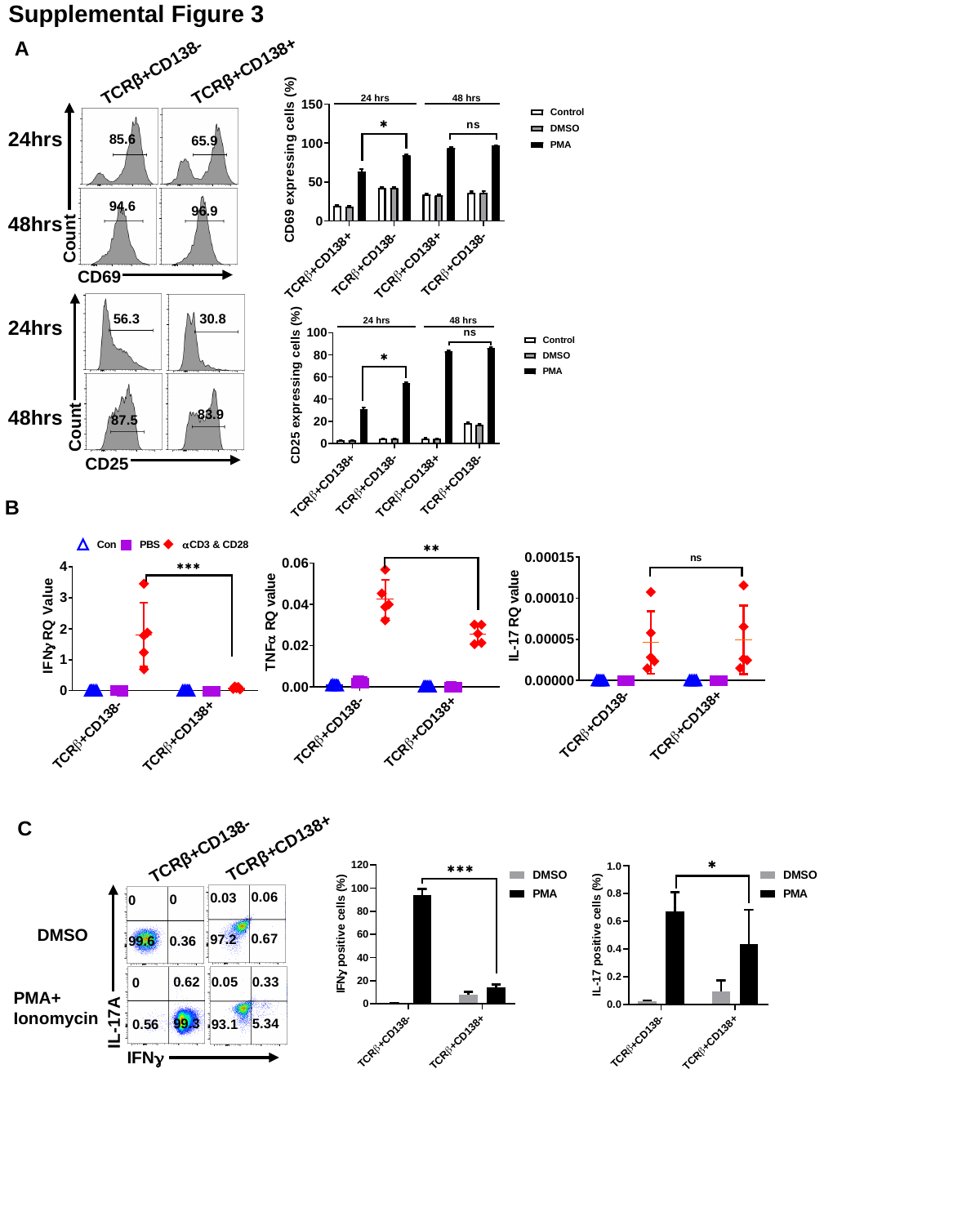

Supplemental Figure 3
A
TCRβ+CD138+
TCRβ+CD138-
24hrs
85.6
65.9
94.6
96.9
48hrs
Count
CD69
30.8
56.3
24hrs
48hrs
83.9
87.5
Count
CD25
B
C
TCRβ+CD138+
TCRβ+CD138-
0.06
0.03
0
0
DMSO
0.67
97.2
0.36
99.6
0.05
0.62
0.33
0
PMA+
Ionomycin
IL-17A
99.3
5.34
0.56
93.1
IFNγ

### Slide 4
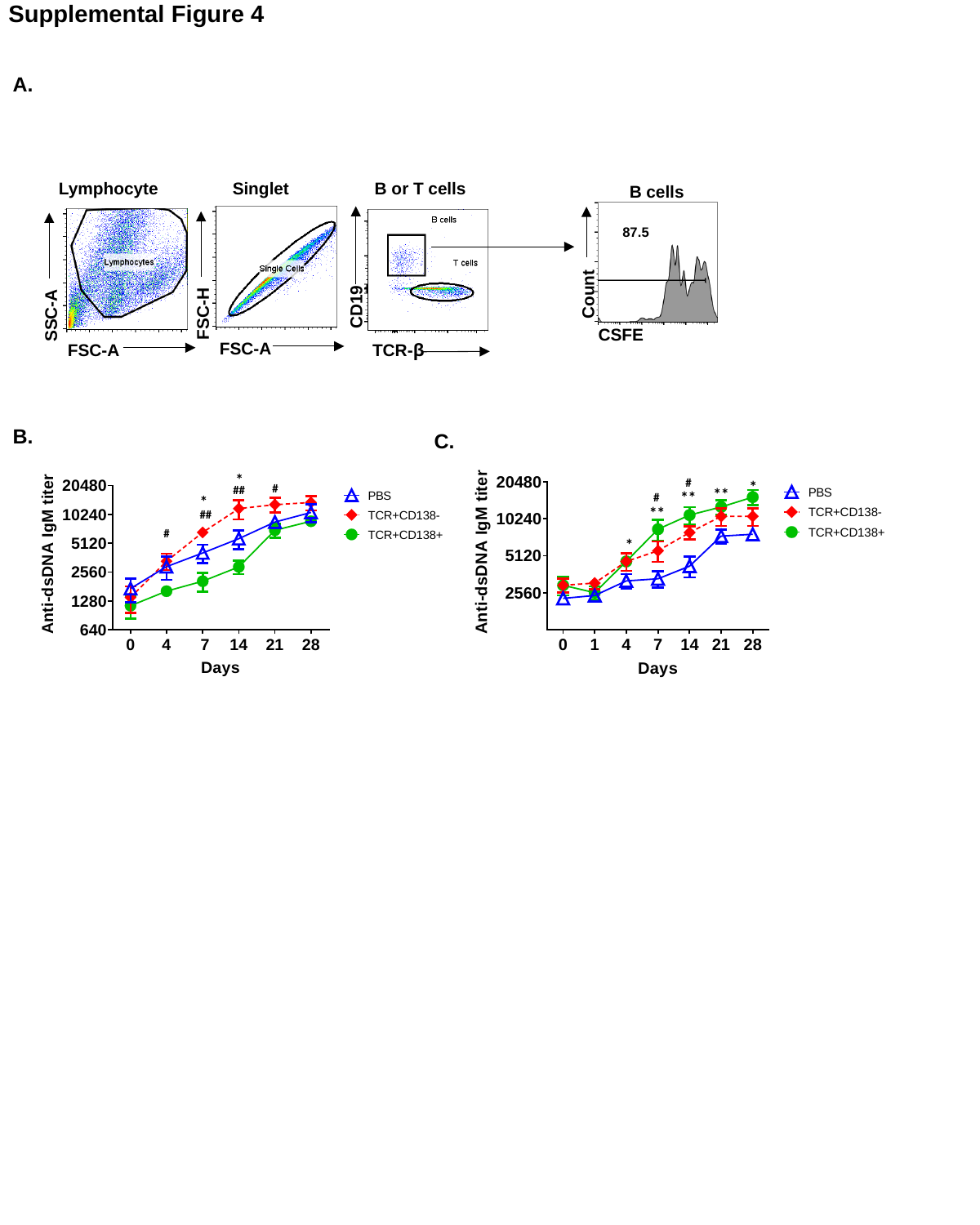

Supplemental Figure 4
A.
Lymphocyte
Singlet
B or T cells
B cells
87.5
Count
CD19
FSC-H
SSC-A
CSFE
FSC-A
FSC-A
TCR-β
B.
C.

### Slide 5
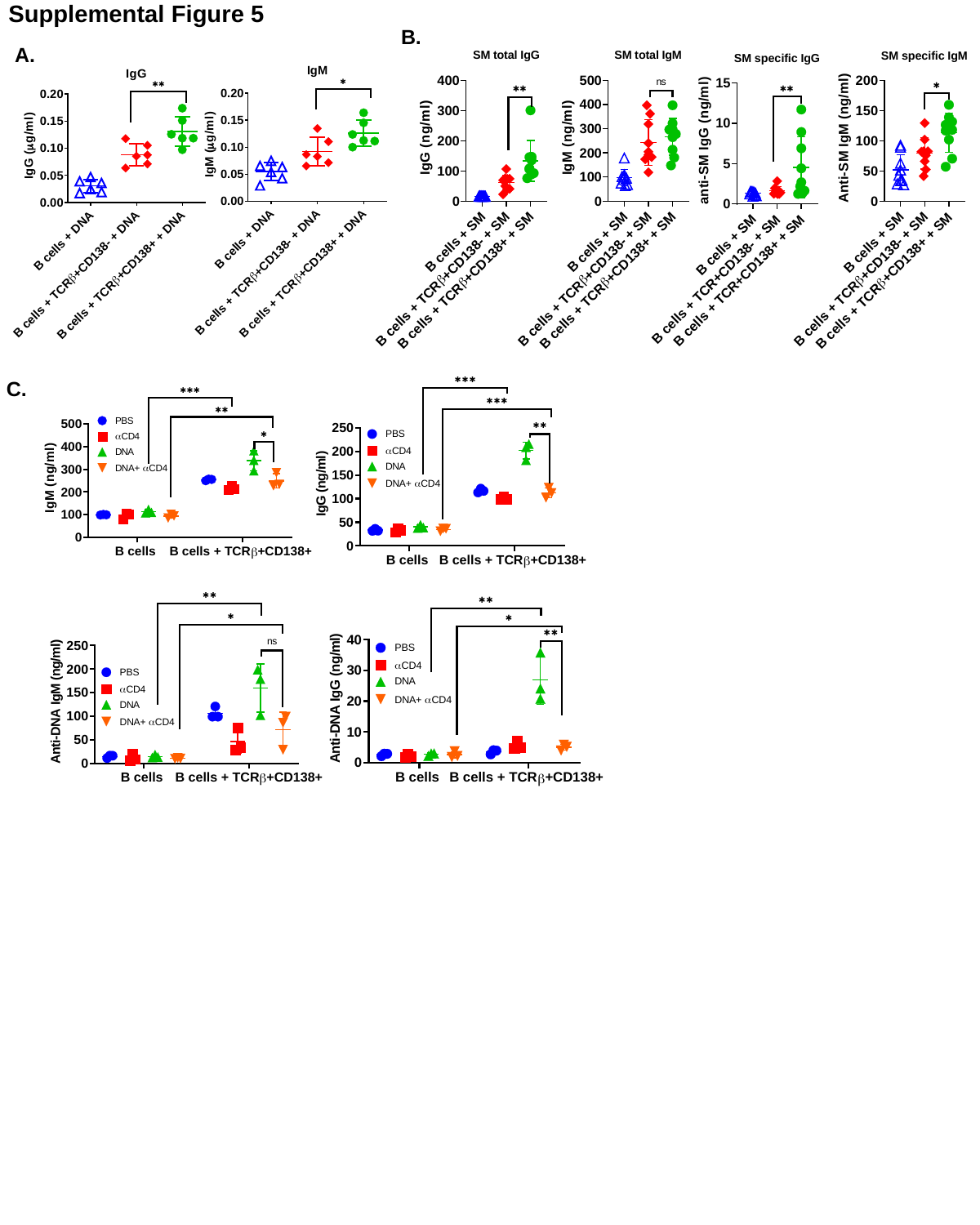

Supplemental Figure 5
B.
A.
C.

### Slide 6
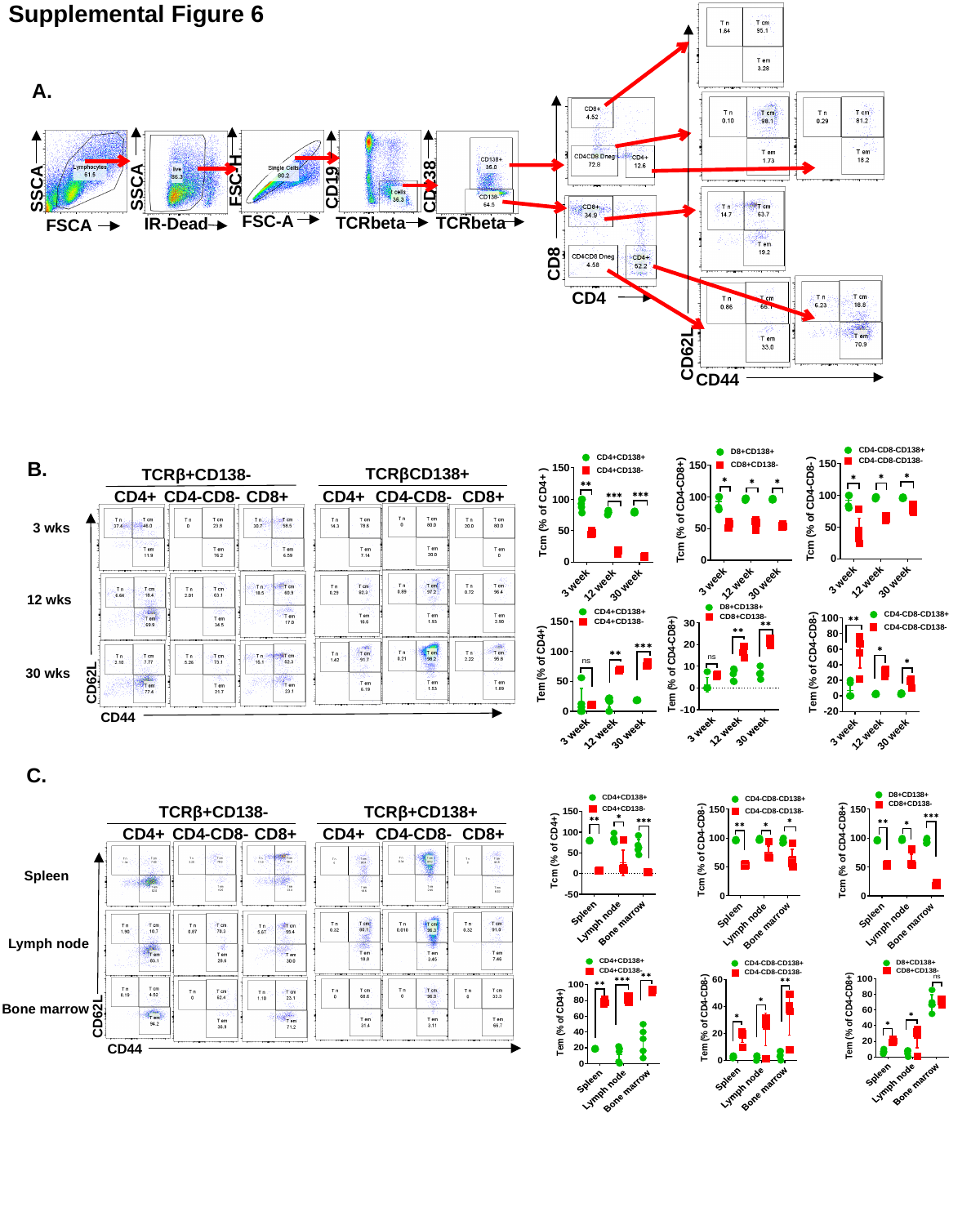

Supplemental Figure 6
A.
FSC-H
SSCA
CD138
CD19
SSCA
FSC-A
TCRbeta
TCRbeta
IR-Dead
FSCA
CD8
CD4
CD62L
CD44
B.
TCRβCD138+
TCRβ+CD138-
CD4+
CD4-CD8-
CD8+
CD4+
CD4-CD8-
CD8+
3 wks
12 wks
30 wks
CD62L
CD44
C.
TCRβ+CD138-
TCRβ+CD138+
CD4+
CD4-CD8-
CD8+
CD4+
CD4-CD8-
CD8+
Spleen
Lymph node
Bone marrow
CD62L
CD44

### Slide 7
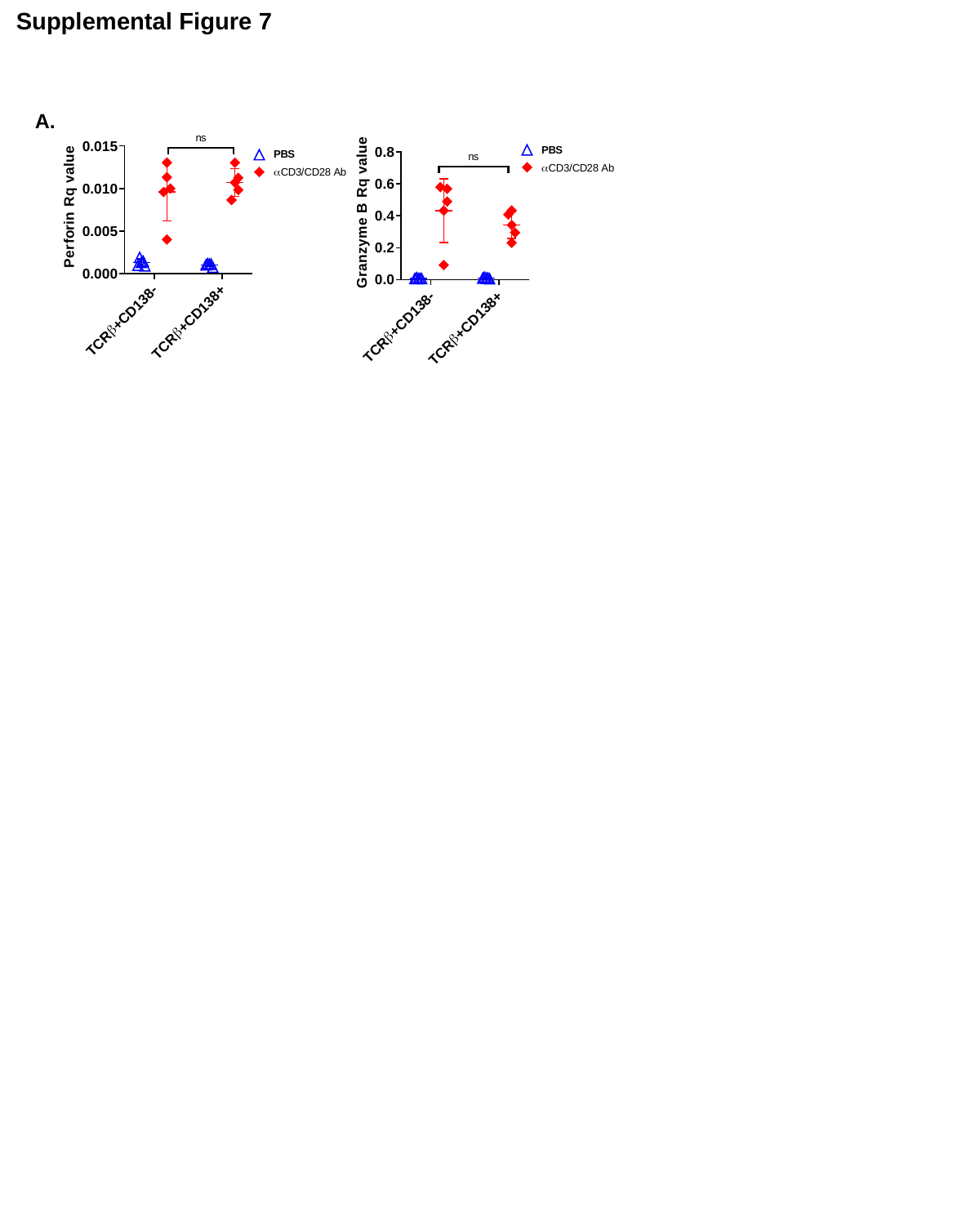

Supplemental Figure 7
A.
